# FCHo2, instead of talin, enables inside-out activation of integrin αvβ5 in curved adhesions

**DOI:** 10.1101/2025.11.28.691229

**Authors:** Chih-Hao Lu, Christina E. Lee, Wei Zhang, Yang Yang, Luis A. Valencia, He You, Ching-Ting Tsai, Bianxiao Cui

## Abstract

Inside-out activation of integrins is crucial for transducing mechanical forces through the extracellular ligand-integrin-talin-F-actin axis. Extensive studies have shown that talin is the essential player in this process by binding to the intracellular tail of β integrins. Here, we show that, while talin binding is essential for inside-out integrin activation in focal adhesions, it is dispensable in curved adhesions - a distinct adhesion architecture that is exclusively mediated by integrin αvβ5 and selectively formed at curved membranes. Instead, a curvature-sensing protein FCHo2 binds to the HDRRE motif in the cytoplasmic tail of integrin β5 (ITGβ5) and inside-out activates integrin αvβ5 in curved adhesions. Intriguingly, FCHo2 does not bind to a similar motif in the homologous integrin β3 tail. Through truncations and mutations, we identified a pivotal tryptophan (W) in the β3 tail, which is conserved in all homologous integrin β isoforms except β5, where it is replaced by a tyrosine (Y766). This tyrosine substitution is crucial for integrin β5’s unique capability in forming curved adhesions. A Y766W mutation abolishes integrin β5’s capacity to form curved adhesions, without affecting its ability to form focal adhesions. Furthermore, our studies suggest that the phosphorylation state of Y766 regulates whether integrin αvβ5 forms curved adhesions or focal adhesions, providing a cellular mechanism governing different adhesion types. Overall, our work unveils distinct molecular interactions and regulatory mechanisms between curved adhesions and focal adhesions, and establishes a molecular basis for the formation of curved adhesions by integrin αvβ5.

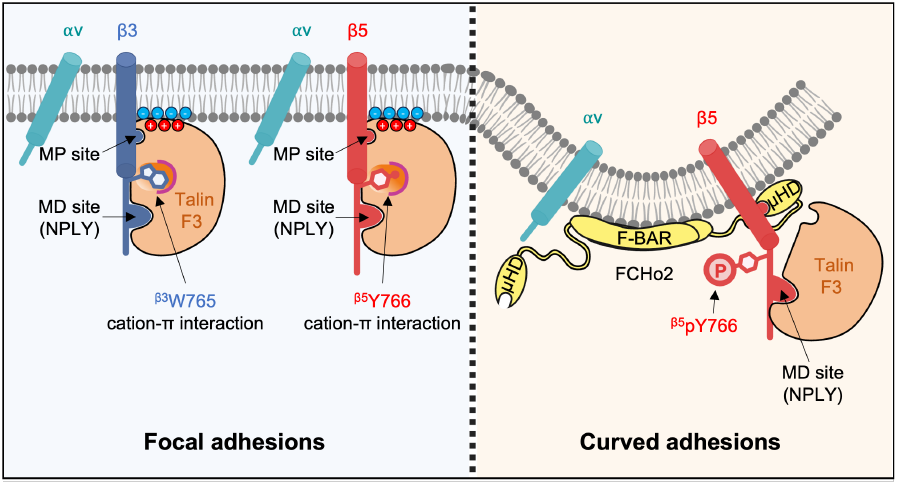

## Introduction

Cells sense their extracellular microenvironments and respond to mechanical cues primarily via integrin-mediated adhesions^1–4^, which confer vital signals to regulate adhesion, migration, differentiation, and proliferation^3,5^. In humans, 18 α and 8 β subunits form 24 heterodimeric integrin complexes that interact with their selective extracellular matrix (ECM) ligands^6–8^. Integrin-mediated adhesions, such as focal adhesions, are regulated bidirectionally, through the intra-cellular binding of adaptor proteins (inside-out signaling) and via the extracellular engagement of ECM ligands (out-side-in signaling)^4,9–12^. Although early studies proposed that inside-out signaling initiates a conformational change in the αβ heterodimer and prime integrins for high-affinity ECM ligand binding^11,13–16^, a recent work demonstrates the opposite sequence: extracellular ligand binding (outside-in signaling) first introduces integrin complex opening, which is subsequently stabilized by inside-out processes that apply tensile forces through the extracellular ligand-integrin-talin-F-actin cytoskeleton axis^12^. In this context, “inside-out integrin activation” refers specifically to intracellular events that stabilize the ligand-induced extended, open conformation of integrins, leading to full activation. Among various cytoplasmic adaptor proteins, talin binding to integrin β intracellular tails is considered an essential and committed step in inside-out activation^9,17–20^. Other cytoplasmic proteins, such as kindlin and paxillin^21–25^, can synergistically enhance integrin activation in cooperation with talin, but cannot trigger inside-out activation independently.

Two decades of extensive studies reveal that talin is involved in inside-out integrin activation through several key interactions. The integrin β tail recruits talin through the high-affinity membrane-distal (MD) NPxY region, which further enables the talin head domain to engage the nearby low-affinity membrane-proximal (MP) region^13,19,26–29^. Talin binding to the MP region disrupts the inhibitory αβ salt bridge and positions itself in a close proximity to the plasma membrane, allowing electrostatic interactions with negatively charged phospholipids^19,28,30,31^. Talin mutations that disrupt the MP or lipid interactions do not affect its overall binding affinity to the integrin β tail *in vitro*, yet strongly inhibit integrin activation in cells^26^. Extensive studies have revealed that talin’s interactions with the MP region and the lipid membrane are both necessary for inside-out activation of integrins^13,26,28,31^.

Upon extracellular ligand engagement, talin also plays a crucial role in the outside-in signaling. Talin directly links to actin filaments to transduce mechanical forces^2,4,32–34^. Furthermore, talin dimerizes and recruits other adaptor proteins such as vinculin, paxillin, and phosphorylated focal adhesion kinase (pFAK) to form large protein assemblies that connect ECM to actomyosin machinery in focal adhesions^2,35–38^. Besides its structural roles in both inside-out and outside-in signalings, talin is the most prominent mechanosensor in integrin-dependent adhesions^20,23^. The autoinhibited globular talin can be activated and stretched to up to 100 nm in length in focal adhesions, exposing 11 cryptic binding sites for vinculin^39^. A talin-based tension sensor shows that talin endures a stretching force well above 10 pN in focal adhesions^40,41^. Force-dependent activation of talin is a key regulatory step in the formation of focal adhesions, requiring a substrate rigidity >5 kPa^42^. As a result, focal adhesions form plentifully on rigid substrates but are sparse in soft environments^43,44^. In natural ECM, collagen forms a fibrous network together with other ECM components, including vitronectin and fibronectin. Most ECM fibers are thin and soft with a bending rigidity below 5 kPa^45^. This aligns with the observation that focal adhesions are sporadic or absent under physiological conditions.

An interesting feature of natural ECM is the cylindrical shape of ECM fibers, which can imprint local membrane curvatures on cell membranes^46–48^ and elicit curvature-dependent signaling events^47,49,50^. Very recently, we identified a distinct integrin-mediated adhesion architecture that exclusively forms at curved plasma membranes, termed “curved adhesions”^40^. Curved adhesions form abundantly in soft 3D fibrous matrices and are molecularly different from focal adhesions^51^ and clathrin-containing adhesions^52,53^. A curvature-sensing protein FCHo2, an essential component of curved adhesions but entirely absent in focal adhesions, serves as a molecular bridge between integrin β5 intracellular domain and membrane curvature. FCHo2 senses and binds to curved membranes through its N-terminal F-BAR domain, and interacts with the intracellular juxtamembrane segment of integrin β5 via its C-terminal microhomology domain (µHD) to convey curvature sensitivity^40^. Talin involves and transmits mechanical forces in both curved adhesions and focal adhesions. However, talin-based tension sensors unveil that talin is under low mechanical tension (3-5 pN) in curved adhesions, but high mechanical tension (>10 pN) in focal adhesions^40^. An intriguing observation is that curved adhesions are exclusively formed by integrin αvβ5, unlike focal adhesions that can be formed by αvβ5, its homolog αvβ3, and most integrin heterodimers^40^. Why curved adhesions and focal adhesions exhibit such distinctive behaviors remains elusive.

In this study, we uncover distinct sets of molecular interactions involved in curved adhesions and focal adhesions. We identify a pivotal and highly conserved tryptophan (W) in integrin β3 and other β isoforms except integrin β5, where W is replaced by a tyrosine (Y766). The W-to-Y substitution in the ITGβ5 tail is crucial for ITGβ5’s unique capability in forming curved adhesions. Our findings suggest that the phosphorylation state of Y766 modulates the equilibrium between focal adhesions and curved adhesions. Furthermore, we find that, although talin binding is crucial for inside-out integrin activation in focal adhesions, it is dispensable in curved adhesions. In curved adhesions, FCHo2 engages the HDRRE motif and inside-out activates integrin αvβ5 independent of talin. From these results, we propose a model that, in focal adhesions, talin engages both the MD and the MP sites, whereas in curved adhesions, talin engages only the MD site, allowing FCHo2 to bind to the HDRRE motif and stabilize active integrin αvβ5 at curved membranes.

## Results

## Talin binding prevents the intracellular domain of ITGβ3, but not that of ITGβ5, from responding to membrane curvature

To induce curved adhesions, we employed a previously-developed vertical nanobar platform which imprints well-defined membrane curvature on the plasma membrane^54–59^. The engineered quartz nanobar arrays were fabricated via photolithography and anisotropic etching on quartz substrates (**Figure 1A**, 200 nm in width, 2 µm in length, 1 µm in height, and 5 µm in spacing). When cells are cultured on the vertically aligned nanobar arrays, cell membranes wrap around nanobars, as confirmed previously by electron microscopy studies^57,60^. For two-dimension fluorescence imaging, the focal plane was positioned at the middle height of nanobars (**Figure 1B**)^55^. Due to the 3D-to-2D projection, the observed curvature effect is primarily located at the half-cylindrically shaped nanobar ends, with a minimal contribution from the top. Substrates were first coated with poly-Llysine (PLL), followed by a crosslinker glutaraldehyde (GA) and then ECM proteins such as vitronectin, fibronectin, or gelatin (**Figure 1B**). Unless specified otherwise, the sub-strates were coated with vitronectin, a high-affinity ECM lig- and for integrin αvβ5. Each quartz substrate contains nanobar areas interlaced with flat areas for direct comparison of cell behaviors in the same culture.

**Figure 1.**
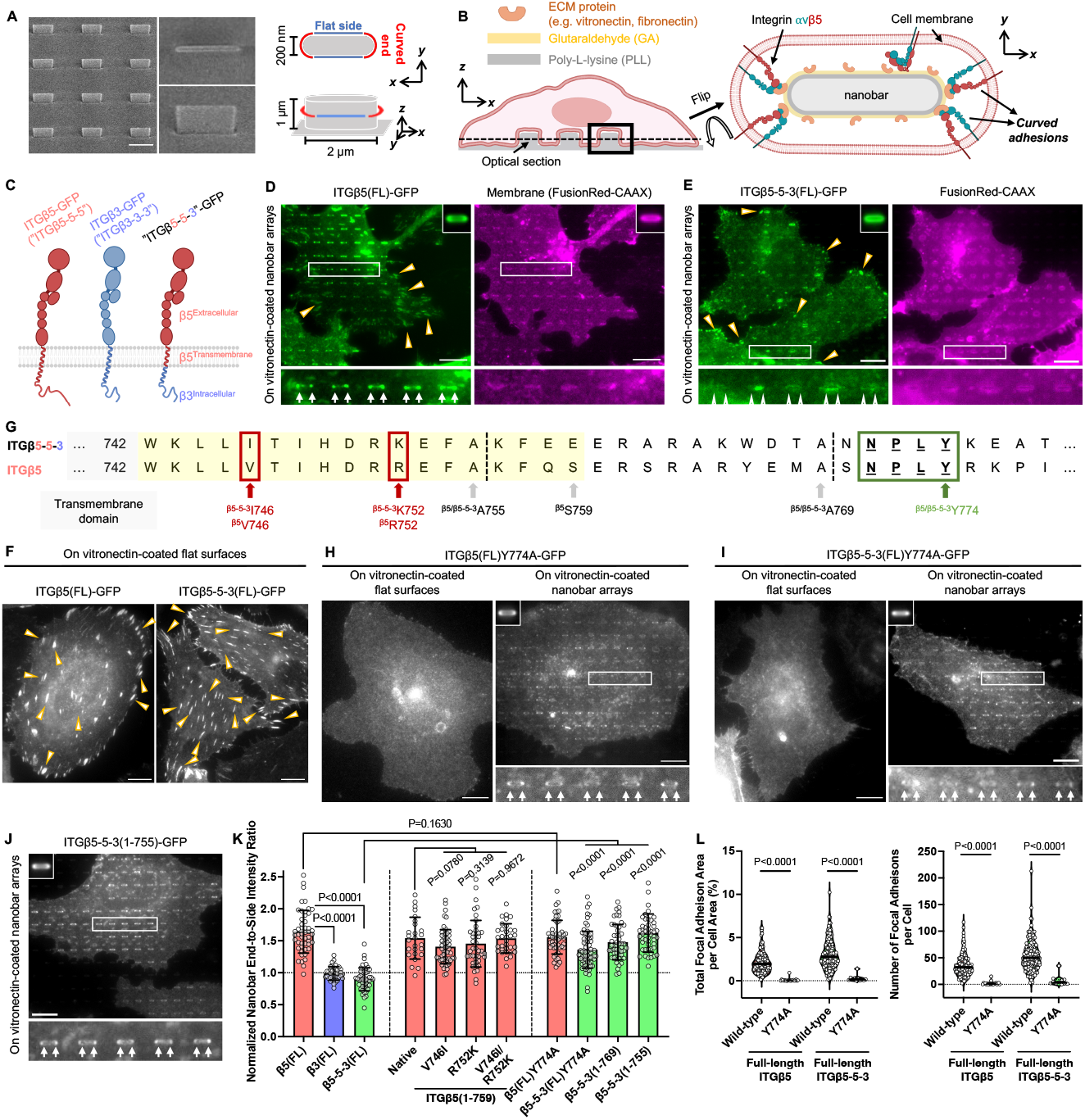
Talin binding prevents the intracellular domain of ITGβ3, but not that of ITGβ5, from responding to membrane curvature. **(A)** Scanning electron microscopic (SEM) images of a nanobar array, and schematic illustration of a single nanobar which induces high curvature at nanobar ends and flat/zero curvature along nanobar sidewalls. All nanobars are 200 nm in width, 2 µm in length, 1 µm in height, and 5 µm in spacing. Scale bar: 2.5 µm. **(B)** Schematic illustrations of a cell interfacing with a nanobar array coated with ECM ligands and formation of curved adhesions. **(C)** Cartoon illustrations of the full-length ITGβ5, ITGβ3, and the ITGβ5-5-3 chimera. **(D)** The full-length ITGβ5-GFP (ITGβ5(FL)-GFP) forms focal adhesions on flat areas between nanobars (yellow arrowheads) and accumulates in curved adhesions formed at nanobar ends (white arrows). The co-expressed membrane marker, FusionRed-CAAX, shows that plasma membranes wrap evenly around nanobars. **(E)** On vitronectin-coated nanobar arrays, ITGβ5-5-3-GFP forms extensive focal adhesions on flat areas between nanobars (yellow arrowheads) but does not accumulate at nanobar ends. **(F)** Both ITGβ5-GFP and ITGβ5-5-3-GFP form abundant focal adhesion architectures on flat substrates (yellow arrowheads). **(G)** Sequence alignment of the intracellular juxtamembrane segments of ITGβ5 and ITGβ5-5-3. The membrane-proximal regions are highlighted by a yellow shade. The two-amino-acid differences are indicated by red rectangular boxes. The high-affinity talin-binding NPLY motifs are denoted by a green rectangular box. **(H)** ITGβ5(Y774A)-GFP, with a point mutation that abolishes talin binding, is unable to form focal adhesions on flat substrates (left), while it shows a clear curvature preference toward nanobar ends on nanobar substrates (right, white arrows). **(I)** ITGβ5-5-3(Y774A)-GFP fails to form focal adhesions on flat substrates (left) but preferentially accumulates at curved membranes induced by nanobar ends (right, white arrows). **(J)** Truncated chimera ITGβ5-5-3(1-755)-GFP preferentially accumulates at nanobar ends (white arrows). **(K)** Quantifications of the nanobar end-to-side intensity ratios, normalized against the ratio of membrane marker FusionRed-CAAX, of ITGβ5(FL)-GFP, ITGβ3(FL)-GFP, ITGβ5-5-3(FL)-GFP, ITGβ5(1-759)-GFP and its mutants, Y774A mutants of ITGβ5(FL)-GFP and ITGβ5-5-3(FL)-GFP, and two ITGβ5-5-3-GFP truncation variants. The dashed line indicates a value of 1. **(L)** Quantifications of the focal adhesion area percentage (left) and the number of focal adhesions per cell (right) of ITGβ5-GFP, ITGβ5-5-3-GFP, and their Y774A mutants. Scale bar: 10 µm for all the cell images. White arrows indicate enrichments at nanobar ends, white empty arrowheads indicate no preferential enrichment at nanobar ends, while yellow arrowheads indicate focal adhesions. In (D)-(E) and (H)-(J), the averaged nanobar images are shown on either the top-right or the top-left corners. Welch’s t tests (unpaired, two-tailed, not assuming equal variance) are applied for statistical analyses of curvature-related measurements (i.e. normalized nanobar end-to-side intensity ratios), while Kolmogorov-Smirnov test was used for statistical analyses of focal adhesions. Error bars represent standard deviations. Each data point represents one cell.

ITGβ5 and ITGβ3 are highly homologous in their amino acid sequences and structures, and both heterodimerize with ITGαv^61,62^. Intriguingly, only ITGβ5, but not ITGβ3 or other integrin β isoforms, responds to membrane curvature and forms curved adhesions^40^. To explore the underlying mechanism, we constructed a green fluorescence protein (GFP)-tagged chimera which consists of the extracellular and trans-membrane domains of ITGβ5, and the intracellular domain of ITGβ3 (β5^Ex^-β5™-β3^In^, named “ITGβ5-5-3”) (**Figure 1C**). Using flat and nanobar substrates, we assessed the capabilities of GFP-tagged, full-length (FL) ITGβ5, ITGβ3, and ITGβ5-5-3 in forming focal adhesions and curved adhesions. In these experiments, vitronectin coating was used for studying ITGβ5 and ITGβ5-5-3, while both fibronectin and vitronectin coatings were used for ITGβ3. On flat areas, both ITGβ5 and ITGβ5-5-3 form extensive focal adhesion patches (**Figure 1F** and **Supplementary Figure 1A**, yellow arrowheads), confirming that ITGβ5-5-3 is a functional integrin. ITGβ3 forms fewer focal adhesions than ITGβ5 and ITGβ5-5-3 (Supplementary Figure 1A, yellow arrowheads), agreeing with the former observations that integrin αvβ3 is less activated than αvβ5 on the cell surface^63^. On nanobar substrates, ITGβ5-GFP preferentially accumulates in curved adhesions formed at nanobar ends (**Figure 1D** and **Supplementary Figure 1B**). In strong contrast, ITGβ5-5-3 does not form curved adhesions and wraps around nanobars with no preferential accumulation at nanobar ends, similar to ITGβ3 (**Figure 1E** and **Supplementary Figure 1B**, white empty triangles). We quantified the curvature preference of ITGβ5, ITGβ3, and ITGβ5-5-3, by calculating their nanobar end-to-side intensity ratios, normalized against the ratio of the membrane marker (FusionRed-CAAX) to account for occasional uneven membrane wrapping (**Supplementary Figure 1C**). Quantifications show the normalized nanobar end-to-side ratio ∼1.6 for ITGβ5, ∼1 for ITGβ3, and ∼0.9 for ITGβ5-5-3 (**Figure 1K**). The end-to-side ratio for ITGβ5-5-3 is less than 1 due to relatively prominent enrichments along the nanobar sidewalls (**Figure 1E**, and **Supplementary Figure 1B**). ITGβ3 exhibits no curvature preference on either fibronectin- or vitronectin-coated nanobar arrays (**Supplementary Figures 1B**; Quantifications in **Supplementary Figure 1D**). Furthermore, when treated with divalent manganese (Mn^2+^), a strong integrin activator, ITGβ3 and ITGβ5-5-3 form extensive focal adhesions (**Supplementary Figure 2A**; Quantifications in **Supplementary Figure 2B**) but remain not enriched at nanobar ends (**Supplementary Figure 2C**; Quantifications in **Supplementary Figure 2D**), indicating that their curvature insensitivities are not due to their insufficient activations on the cell surface.

**Figure 2.**
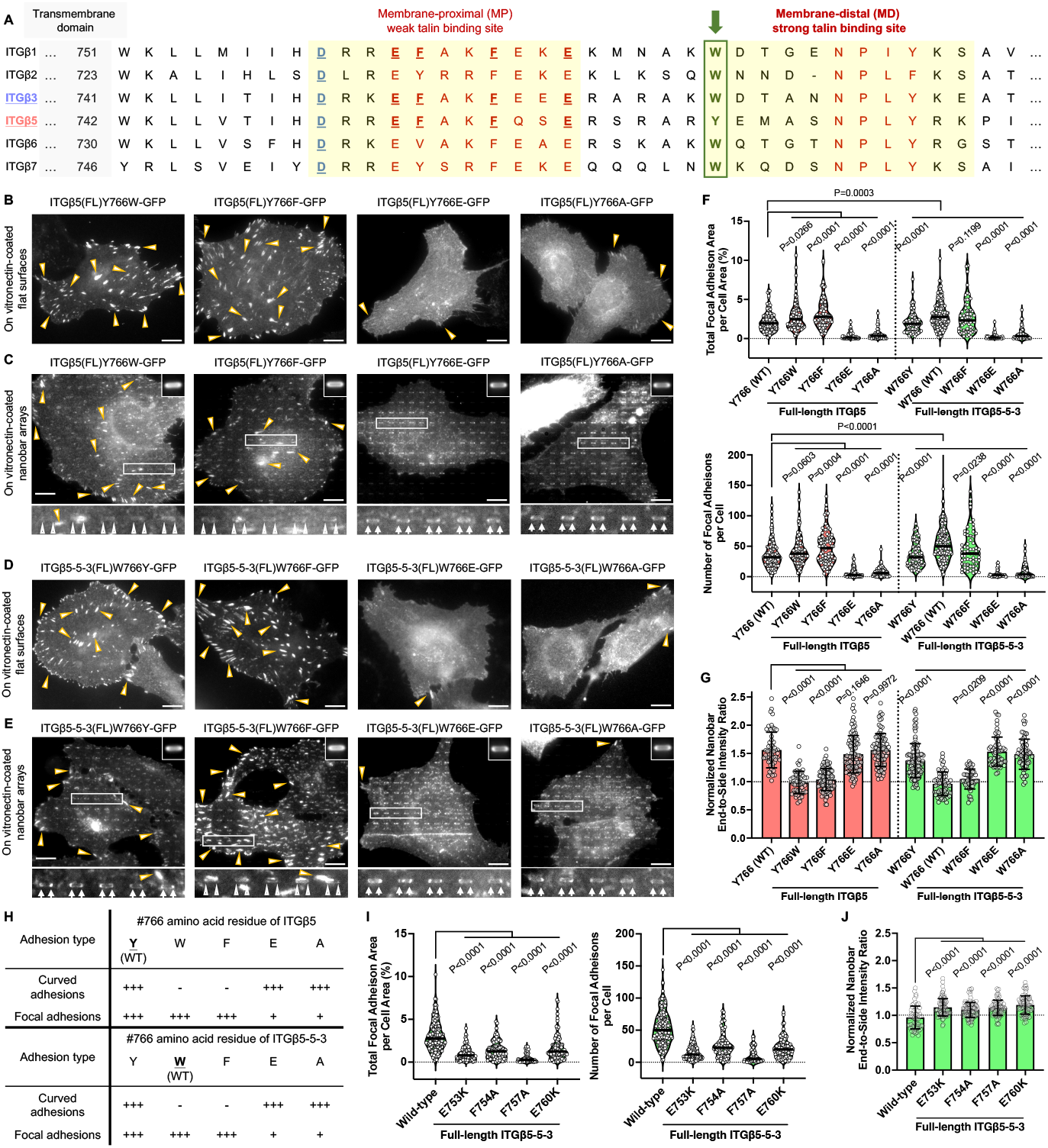
The tryptophan-to-tyrosine substitution at the pivotal location is crucial for ITGβ5’s unique capability in forming curved adhesions. **(A)** Sequence alignments of the intracellular domains of homologous integrin β isoforms. Two talin binding sites (membrane-proximal (MP) and membrane-distal (MD) regions) are highlighted by yellow shades. The amino acid residues identified to be essential for membrane-proximal talin binding are red-colored, bolded, and underlined. The salt bridge-forming aspartate residues are blue-colored, bolded and underlined. The pivotal tryptophan/tyrosine residues are denoted by a green rectangular box. **(B)** ITGβ5(Y766W)-GFP and ITGβ5(Y766F)-GFP form considerable focal adhesions on flat substrates (yellow arrowheads), while there are only few focal adhesion patches formed by ITGβ5(Y766A)-GFP or ITGβ5(Y766E)-GFP. **(C)** On nanobars, both ITGβ5(Y766A)-GFP and ITGβ5(Y766E)-GFP display clear preferences for nanobar ends (white arrows), while neither ITGβ5(Y766W)-GFP nor ITGβ5(Y766F)-GFP shows an apparent curvature preference on nanobar substrates. **(D)** ITGβ5-5-3(W766Y)-GFP and ITGβ5-5-3(W766F)-GFP form extensive focal adhesions on flat substrates (yellow arrowheads), while there are only few focal adhesions formed by ITGβ5-5-3(W766A)-GFP or ITGβ5-5-3(W766E)-GFP. **(E)** On nanobars, ITGβ5-5-3(W766Y)-GFP, ITGβ5-5-3(W766A)-GFP, and ITGβ5-5-3(W766E)-GFP all preferentially accumulate at nanobar ends (white arrows), while ITGβ5-5-3(Y766F)-GFP shows not curvature preference. **(F)** Quantifications of the focal adhesion area percentage (top) and the number of focal adhesions per cell (bottom) of ITGβ5-GFP, ITGβ5-5-3-GFP, and their Y766/W766 mutants. **(G)** Quantifications of the normalized nanobar end-to-side intensity ratio of ITGβ5-GFP, ITGβ5-5-3-GFP, and their Y766/W766 mutants. **(H)** A table summarizing curved adhesion- and focal adhesion-forming capability of ITGβ5, ITGβ5-5-3, and their Y766/W766 mutants. **(I)** Quantifications of the focal adhesion area percentage (left) and the number of focal adhesions per cell (right) of ITGβ5-5-3-GFP and its four variants with key mutations in the MP talin-binding site. **(J)** Quantifications of the normalized nanobar end-to-side intensity ratio of ITGβ5-5-3-GFP and its MP talin-binding mutants. Scale bar: 10 µm for all the cell images. White arrows indicate enrichments at nanobar ends, white empty arrowheads indicate no preferential enrichment at nanobar ends, while yellow arrowheads indicate focal adhesions. In (C) and (E), the averaged nanobar images are shown on the top-right corners. In (G) and (J), the dashed line indicates a value of 1. Welch’s t tests (unpaired, two-tailed, not assuming equal variance) are applied for statistical analyses of curvature-related measurements, while Kolmogorov-Smirnov test was used for statistical analyses of focal adhesions. Error bars represent standard deviations. Each data point represents one cell.

The intracellular juxtamembrane region of ITGβ5 is crucial for its interaction with FCHo2 and its curvature preference^40^. ITGβ5-5-3 and ITGβ5 differ by only two amino acids (^β5^V746 versus ^β5-5-3^I746, and ^β5^R752 versus ^β5-5-3^K752) in their juxtamembrane regions (**Figure 1G**, denoted by red rectangular boxes), To determine whether these two amino acids account for the distinctive curvature preferences between ITGβ5 and and ITGβ5-5-3, we truncated ITGβ5 from its C-terminus into ITGβ5(1-759), which contains the critical juxtamembrane region (**Figure 1F**, highlighted by a yellow shade) and has reportedly exhibited a strong curvature preference^40^. We subsequently constructed two single mutants-ITGβ5(1-759)V746I and ITGβ5(1-759)R752K, and a double mutant-ITGβ5(1-759)V746I/R752K. Surprisingly, both single and double mutants display clear curvature preferences, resembling the native ITGβ5(1-759) (**Supplementary Figure 1E**, white arrows; Quantifications in **Figure 1K**). These results indicate that the disparate curvature preferences between ITGβ5 and ITGβ5-5-3 do not arise from their two-amino-acid differences in the juxtamembrane region.

Based on these results, we hypothesized that elements in the membrane-distal region of ITGβ3 intracellular tail likely inhibit ITGβ5-5-3 from responding to membrane curvature. In the membrane-distal region of ITGβ3 intracellular tail, the NPLY motif serves as the high-affinity binding site for talin (**Figure 1G**; highlighted by a green rectangular box). To determine whether talin binding impedes ITGβ5-5-3 from sensing membrane curvature, we engineered the full-length ITGβ5(Y774A)-GFP and ITGβ5-5-3(Y774A)-GFP, with a Y-to-A mutation in the NPLY motif known to abrogate talin-ITGβ interactions^29,64–66^. Indeed, without talin engagement, both ITGβ5(Y774A)-GFP and ITGβ5-5-3(FL)Y774A fail to form focal adhesions on flat substrates (**Figure 1H-1I**; Quantifications in **Figure 1L**), consistent with previous reports^14,67^. Surprisingly, both mutants demonstrate prominent accumulations at nanobar ends (**Figure 1H-1I**, white arrows; Quantifications in **Figure 1K**). Therefore, the curvature preference of ITGβ5-5-3 can be restored by abolishing talin engagement.

To further test the hypothesis that talin binding to ITGβ5-5-3 cytoplasmic tail prevents it from sensing membrane curvature, we constructed two C-terminal truncation variants of ITGβ5-5-3, with ITGβ5-5-3(1-769) truncated immediately upstream of the NPLY motif and ITGβ5-5-3(1-755) containing only the juxtamembrane segment (**Figure 1G**; denoted by gray arrows). Remarkably, both ITGβ5-5-3(1-755) and ITGβ5-5-3(1-769) show clear preferences for nanobar ends (**Figure 1J** and **Supplementary Figure 1F**, white arrows; Quantifications in **Figure 1K**). These results confirm that the juxtamembrane region of ITGβ3 cytoplasmic tail is capable of responding to membrane curvature, however, talin binding to the membrane-distal region prevents ITGβ5-5-3 from reacting to membrane curvature.

### The pivotal tryptophan-to-tyrosine substitution is crucial for ITGβ5’s unique capability in forming curved adhesions

Now, a puzzle emerges since talin binds to the membrane-distal NPLY motif in both ITGβ5 and ITGβ5-5-3. While talin binding blocks the curvature preference of ITGβ5-5-3, this inhibition surprisingly doesn’t extend to ITGβ5 (i.e. ITGβ5-5-5). To understand this discrepancy, we carefully examined the intracellular domain sequence of homologous integrin β isoforms (**Figure 2A**). A noteworthy observation comes to light -a pivotal tryptophan (W) located upstream of the NPLY talin-binding region is conserved in all the integrin β isoforms except ITGβ5, in which the tryptophan is replaced by a tyrosine (Y) (**Figure 2A**, highlighted by a green rectangular box). From the crystal structures, the pivotal tryptophan (^β3^W765) of ITGβ3 is deeply wedged within the pocket of talin-1 F3 domain, with the indole ring of the pivotal tryptophan positioned parallel to the cationic guanidinium group of the arginine (^talin-1^R358) within the F3 domain, resulting in the formation of a significant cation-π interaction^13,26^. Although ^β5^Y766 is expected to maintain a similar interaction, phosphorylation of ^β5^Y766 results in a bulky, negatively charged phosphotyrosine that does not fit into the pocket of talin-1 F3 and potentially alters such interactions. We hypothesized that the W-to-Y substitution is crucial for the formation of ITGβ5-mediated curved adhesion.

To scrutinize this hypothesis, we engineered the full-length ITGβ5(Y766W) by replacing the tyrosine (Y766) with a tryptophan. Strikingly, the Y-to-W mutation completely abolishes the curvature preference, with ITGβ5(Y766W) forming considerable focal adhesions on flat substrates (**Figure 2B**, yellow arrowheads) but no curved adhesions at nanobar ends (**Figure 2C**, white empty triangles), analogous to ITGβ5-5-3. Conversely, we also engineered a reverse W-to-Y substitution in the full-length ITGβ5-5-3 by replacing the tryptophan (W766) with a tyrosine, resulting in ITGβ5-5-3(W766Y). The W-to-Y mutation surprisingly restores the curvature preference, with ITGβ5-5-3(W766Y) forming both focal adhesions on flat substrates (**Figure 2D**, yellow arrowheads) and curved adhesions at nanobar ends (**Figure 2E**, white arrows). Therefore, ITGβ5-5-3(W766Y) enables the formation of both curved adhesion and focal adhesions, similar to ITGβ5(FL). Jointly, the tryptophan residue in the pivotal location supports only the formation of focal adhesions, but the tyrosine substitution promotes the formation of both curved adhesions and focal adhesion.

Tyrosine residues can be phosphorylated in cells. A previous study suggests that three tyrosine residues in the ITGβ5 intracellular domain, including Y766, can be phosphorylated^68^. Due to the lack of a pY766-specific ITGβ5 antibody, we used phenylalanine (F) to mimic the non-phosphorylatable tyrosine and glutamate (E) as a phosphotyrosine mimetic, to investigate whether Y766 phosphorylation affects the formation of focal adhesions and/or curved adhesions. Additionally, phenylalanine is expected to form the significant cation-π interaction with ^talin-1^R358, while the glutamate cannot. For this study, we engineered four more full-length mutants: ITGβ5(Y766F), ITGβ5(Y766E), ITGβ5-5-3(W766F), and ITGβ5-5-3(W766E). We found that both ITGβ5(Y766F) and ITGβ5-5-3(W766F) form substantial focal adhesions on flat areas (**Figure 2B** and **2D**, yellow arrowheads), but fail to form curved adhesions at nanobar ends (**Figure 2C** and **2E**, white empty triangles). On the contrary, ITGβ5(Y766E) and ITGβ5-5-3(W766E) form only few focal adhesions on flat substrates (**Figure 2B** and **2D**, yellow arrowheads), but they form considerable curved adhesions with strong accumulations at nanobar ends (**Figure 2C** and **2E**, white arrows). The sharp contrasts between ITGβ5(Y766F) and ITGβ5(Y766E), and between ITGβ5-5-3(W766F) and ITGβ5-5-3(W766E), implicate that the phosphorylation state of Y766 regulates whether ITGβ5 can assemble focal adhesions or curved adhesions.

To further understand the role of the cation-π interaction in the formation of focal adhesions and curved adhesions, we mutated the tryptophan (^β5-5-3^W766) and the tyrosine (^β5^Y766) into an alanine (A), which possesses a nonpolar methyl side chain and unable to participate in the cation-π interaction. Very interestingly, on flat surfaces, both ITGβ5-5-3(W766A) and ITGβ5(Y766A) only form scarce focal adhesions (**Figure 2B** and **2D**, yellow arrowheads), while they both display strong curvature preferences on nanobar substrates (**Figure 2C** and **2E**, white arrows), indicating that the cation-π interactions between talin F3 subdomain and ITGβ5/β3 tails enhance the formation of focal adhesions, but at the same time, inhibits the formation of curved adhesions.

We quantified the focal adhesions and curved adhesions for the five variants (Y766, W766, F766, E766, and A766) from 10 constructs (5 for ITGβ5 and 5 for ITGβ5-5-3). Curved adhesions are reflected by the normalized nanobar end-to-side ratios. Among the five variants of both ITGβ5 and ITGβ5-5-3, W766 and F766 exclusively form focal adhesions (**Figure 2F-2G**), while E766 and A766 primarily form curved adhesions (**Figure 2F-2G**). The drastically attenuated focal adhesion formation by A766 and E766 mutations agrees with previous studies proving that the W-to-A and W-to-E mutations of ITGβ1^27^ and the W-to-A mutation of ITGβ3^17,26^ lead to reduced focal adhesion formation and lower affinity for talin, compared with their wild-type counterparts. When talin-1 is co-overexpressed, we observed that both ITGβ5-5-3(W766A) and ITGβ5-5-3(W766E) can form substantial focal adhesions, unlike the ITGβ5-5-3(Y774A) mutant (**Supplementary Figure 3**, yellow arrowheads). Therefore, these mutants can still interact with talin. Among these mutants, only the variants with Y766 can form both focal adhesions and curved adhesions, likely due to its phosphorylation state (**Figure 2H**).

**Figure 3.**
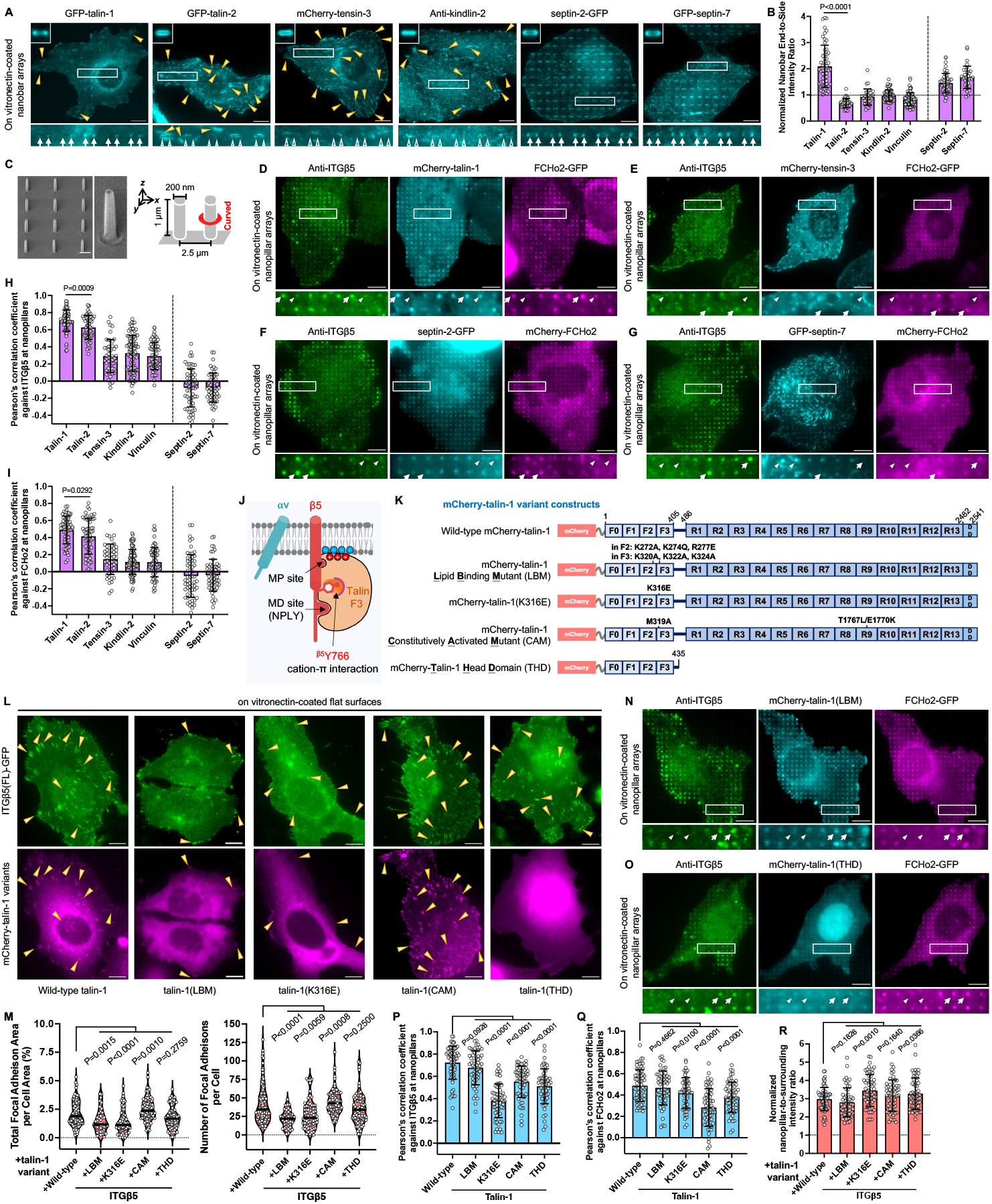
Talin-1, but not kindlin, tensin, or septin, is the primary mechanosensitive component in curved adhesions. **(A)** Representative images of GFP-talin-1, GFP-talin-2, mCherry-tensin-3, anti-kindlin-2, septin-2-GFP, and GFP-septin-7 on nanobar substrates. Talin-1, septin-2, and septin-7 all preferentially cluster at nanobar ends (white arrows). Whereas, talin-2, tensin-3, and kindlin-2 show no preference for curved membranes induced at nanobar ends (white empty arrowheads). Additionally, talin-1, talin-2, tensin-3, and kindlin-2 are all localized to focal adhesion patches formed on the flat regions of nanobar substrates (yellow arrowheads). The averaged nanobar images are shown on the top-left corners. **(B)** Quantifications of the normalized nanobar end-to-side intensity ratios of five cytoskeletal adaptors and septins. **(C)** SEM images of a nanopillar array and schematic illustration of vertical nanopillars inducing membrane curvature by their cylindrical shape. All nanopillars are 200 nm in diameter, 1 µm in height, and 2.5 µm in spacing. Scale bar: 1 µm. **(D-E)** Anti-ITGβ5, FCHo2-GFP, and mCherry-talin-1 accumulate and colocalize in curved adhesions formed at nanopillar locations (D), while mCherry-tensin-3 is not involved in curved adhesions, as seen by their modest-to-low accumulations and low spatial correlations with anti-ITGβ5 and FCHo2-GFP, at nanopillars (E). **(F-G)** Although both septin-2-GFP (F) and GFP-septin-7 (G) preferentially cluster at nanopillars, they are not spatially correlated with anti-ITGβ5 and mCherry-FCHo2. **(H-I)** Quantifications of the Pearson’s correlation coefficients between five cytoskeletal adaptors or septins and ITGβ5 (H), or and FCHo2 (I), at nanopillars. **(J)** Cartoon illustration of talin F3 subdomain interacting with ITGβ5 intracellular tail and plasma membrane. **(K)** Domain structures of five mCherry-talin-1 variants. F: FERM domains; R: rod domains; DD: dimerization domain. **(L)** ITGβ5-GFP forms abundant focal adhesion patches on flat substrates when co-expressed with the wild-type mCherry-talin-1, mCherry-talin-1(CAM), or mCherry-talin-1(THD). However, it forms fewer and smaller focal adhesion structures when either mCherry-talin-1(LBM) or mCherry-talin-1(K316E) is co-expressed. **(M)** Quantifications of the focal adhesion area percentage (left) and the number of focal adhesions per cell (right) of ITGβ5-GFP, when co-expressed with five mCherry-talin-1 variants. **(N)** Anti-ITGβ5, FCHo2-GFP, and mCherry-talin-1(LBM) all accumulate and colocalize in curved adhesions formed at nanopillars. **(O)** mCherry-talin-1(THD) displays low spatial correlations with anti-ITGβ5 and FCHo2-GFP at nanopillars. **(P-Q)** Quantifications of Pearson’s correlation coefficients between anti-ITGβ5 and mCherry-talin-1 variants (P), or between FCHo2-GFP and mCherry-talin-1 variants (Q), at nanopillars. **(R)** Quantifications of the nanopillar-to-surrounding intensity ratios, normalized against the ratio of membrane marker FusionRed-CAAX, of ITGβ5 when co-expressed with various mCherry-talin-1 variants. Scale bar: 10 µm for all the cell images. White arrows indicate enrichments at nanobar ends, white empty arrowheads indicate no preferential enrichment at nanobar ends, while yellow arrowheads indicate focal adhesions. In (D)-(G) and (N)-(Q), white arrows indicate high-intensity correlations, while white triangles indicate low-intensity correlations at nanopillars. In (B) and (R), the dashed line indicates a value of 1. Welch’s t tests (unpaired, two-tailed, not assuming equal variance) are applied for statistical analyses of curvature-related measurements (i.e. normalized nanobar end-to-side intensity ratios and Pearson’s correlation coefficients at nanopillars). Error bars represent standard deviations. Each data point represents one cell.

In the intracellular domains of β integrins, the low-affinity MP binding site for talin overlaps with the juxtamembrane region crucial for FCHo2 binding^40^. To investigate the hypothesis that talin binding to the MP site prevents FCHo2 engagement and thus inhibits the curvature enrichment of ITGβ5-5-3, we introduced mutations into the full-length ITGβ5-5-3 to weaken talin binding to the MP site^13,27,28^, including ITGβ5-5-3(FL)E753K, ITGβ5-5-3(FL)F754A, ITGβ5-5-3(FL)F757A, and ITGβ5-5-3(FL)E760K (**Figure 2A**, red-colored, bolded and underlined). All of these mutants form fewer and smaller focal adhesions than ITGβ5-5-3(FL) (**Supplementary Figure 4A**, yellow arrowheads; Quantifications in **Figure 2I**), in accord with previous studies^13,27,28^. Interestingly, these mutants all show higher curvature enrichments than the wild-type ITGβ5-5-3 (**Supplementary Figure 4B**, white arrows; Quantifications in **Figure 2J**), supporting the hypothesis. Nevertheless, these single mutations do not completely abolish talin binding to the MP site, such that all these mutants still exhibit weaker curvature preferences than ITGβ5-5-3(W766Y/E/A) mutants, ITGβ5-5-3(Y774A), and the ITGβ5-5-3 truncation variants.

**Figure 4.**
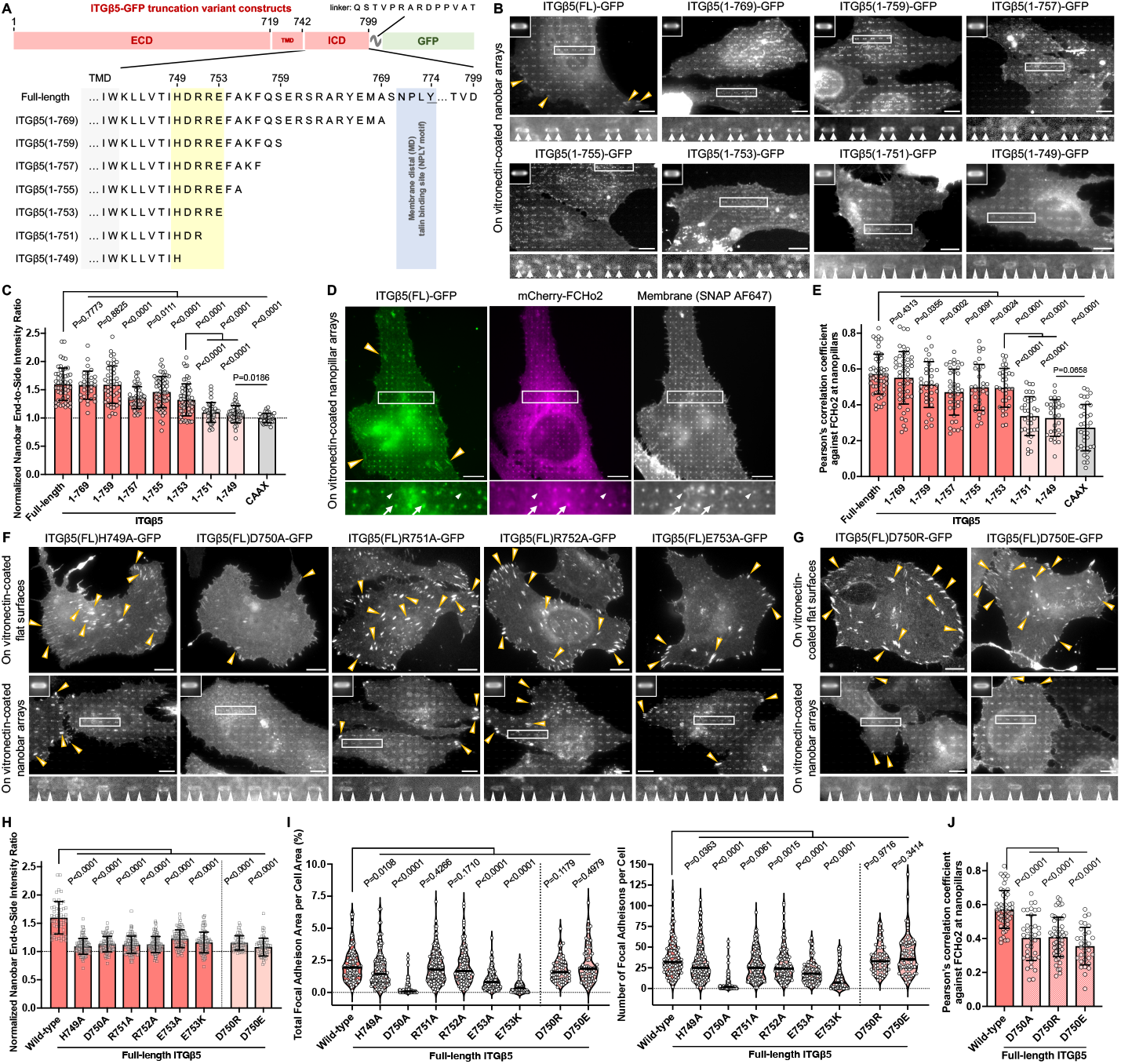
FCHo2 engages the HDRRE motif of ITGβ5 to convey curvature sensitivity. **(A)** Amino acid sequences of the juxtamembrane segment of ITGβ5-GFP and its truncation variants. The HDRRE motif for FCHo2 binding is highlighted by a yellow shade. ECD: extracellular domain; TMD: transmembrane domain; ICD: intracellular domain. **(B)** Representative images of ITGβ5 and its truncation variants on nanobar arrays. resembling the full-length ITGβ5, ITGβ5(1-769), ITGβ5(1-759), ITGβ5(1-757), ITGβ5(1-755), and ITGβ5(1-753) all preferentially enrich at nanobar ends (white arrows). Whereas, ITGβ5(1-751) and ITGβ5(1-749) wrap around nanobars with no preferential accumulation at nanobar ends (white empty arrow-heads). **(C)** Quantifications of the normalized nanobar end-to-side intensity ratios of ITGβ5-GFP truncation variants. **(D)** ITGβ5(FL)-GFP and mCherry-FCHo2 show a strong correlation at individual nanopillars. Cell membranes were visualized via Alexa Fluor 647 SNAP-tag labeling. White arrows indicate high-intensity correlations, while white triangles indicate low-intensity correlations at nanopillars. **(E)** Quantifications of the Pearson’s correlation coefficients between mCherry-FCHo2 and ITGβ5-GFP truncation variants at nanopillars. **(F)** Representative images of the full-length ITGβ5-GFP HDRRE mutants, including ITGβ5(H749A), ITGβ5(D750A), ITGβ5(R751A), ITGβ5(R752A), and ITGβ5(E753A) on flat substrates (top row) or nanobar arrays (bottom row). Most mutations in the HDRRE motif do not significantly affect ITGβ5-mediated focal adhesions (yellow arrow-heads), but all the ITGβ5 HDRRE single mutants demonstrate much reduced curvature preferences for nanobar ends (white empty arrowheads) compared with the full-length ITGβ5. **(G)** Representative images of the full-length ITGβ5-GFP D750 mutants, including ITGβ5(D750R) and ITGβ5(D750E), on flat substrates (top row) or nanobar arrays (bottom row). Both ITGβ5(D750R) and ITGβ5(D750E) form plentiful focal adhesions on flat substrates (yellow arrowheads). However, neither of them forms curved adhesions at nanobar ends (white empty arrowheads). **(H)** Quantifications of the normalized nanobar end-to-side intensity ratios of ITGβ5-GFP and its HDRRE mutants. **(I)** Quantifications of the focal adhesion area percentage (left) and the number of focal adhesions per cell (right) of ITGβ5-GFP and its HDRRE mutants. **(J)** Quantifications of Pearson’s correlation coefficients between mCherry-FCHo2 and ITGβ5-GFP D750 mutants at nanopillars. Scale bar: 10 µm for all the cell images. White arrows indicate enrichments at nanobar ends, white empty arrowheads indicate no preferential enrichment at nanobar ends, while yellow arrowheads indicate focal adhesions. In (B), (F), and (G), the averaged nanobar images are shown on the top-left corners. In (C) and (H), the dashed line indicates a value of 1. Welch’s t tests (unpaired, two-tailed, not assuming equal variance) are applied for statistical analyses of curvature-related measurements, while Kolmogorov-Smirnov test was used for statistical analyses of focal adhesions. Error bars represent standard deviations. Each data point represents one cell.

### Talin-1, but not talin-2, kindlin, tensin, or septins, is the primary mechanosensitive module in curved adhesions

Many intracellular adaptor proteins participate in integrin-mediated force transmission for inside-out integrin activation. Among these adaptors, talin, kindlin, and tensin have been considerably documented. Talin and tensin both bind to the NPxY motif of integrin β subunits (**Figure 2A**), while kindlin interacts with integrin β tails through the NxxY motif downstream of the NPxY motif^22,24,69,70^. We previously showed that curved adhesions involve a subset of focal adhesion proteins, including talin-1, paxillin, and zyxin, but not vinculin and pFAK^40^. Here, we examined whether talin-2 (a highly homologous isoform of talin-1), tensin, and kindlin are involved in curved adhesions. Tensin-3 and kindlin-2 isoforms were chosen owing to their abundance and reported roles in integrin-mediated adhesions^71–79^. Moreover, talin-2 and tensin-3 have also been shown to participate in integrin αvβ5-mediated reticular adhesions formed on flat substrates^53^.

On flat surfaces, talin-1, talin-2, tensin-3, kindlin-2 (endogenous), and vinculin (endogenous) all colocalize with the endogenous ITGβ5 in focal adhesions and/or “adhesion-like” patches, both of which are devoid of FCHo2, the key component of curved adhesions (**Supplementary Figure 5A-5E**). On nanobars, however, only talin-1 displays clear accumulations at nanobar ends, in addition to enrichments in focal adhesions at the cell peripheral (**Figure 3A**, white arrows and yellow arrowheads). Talin-2, tensin-3, kindlin-2, and vinculin show no preference for nanobar ends despite their strong localizations to adhesion architectures on the flat regions between nanobars (**Figure 3A** and **Supplementary Figure 6A**, white empty triangles and yellow arrowheads). Overexpression of talin-2 and tensin-3 significantly enhance the formation of adhesion patches on flat areas (**Supplementary Figure 5B-5C**). Quantifications of the normalized end-to-side ratios confirm that only talin-1, but not talin-2, tensin-3, kindlin-2, or vinculin, clusters at curved membranes (**Figure 3B**).

**Figure 5.**
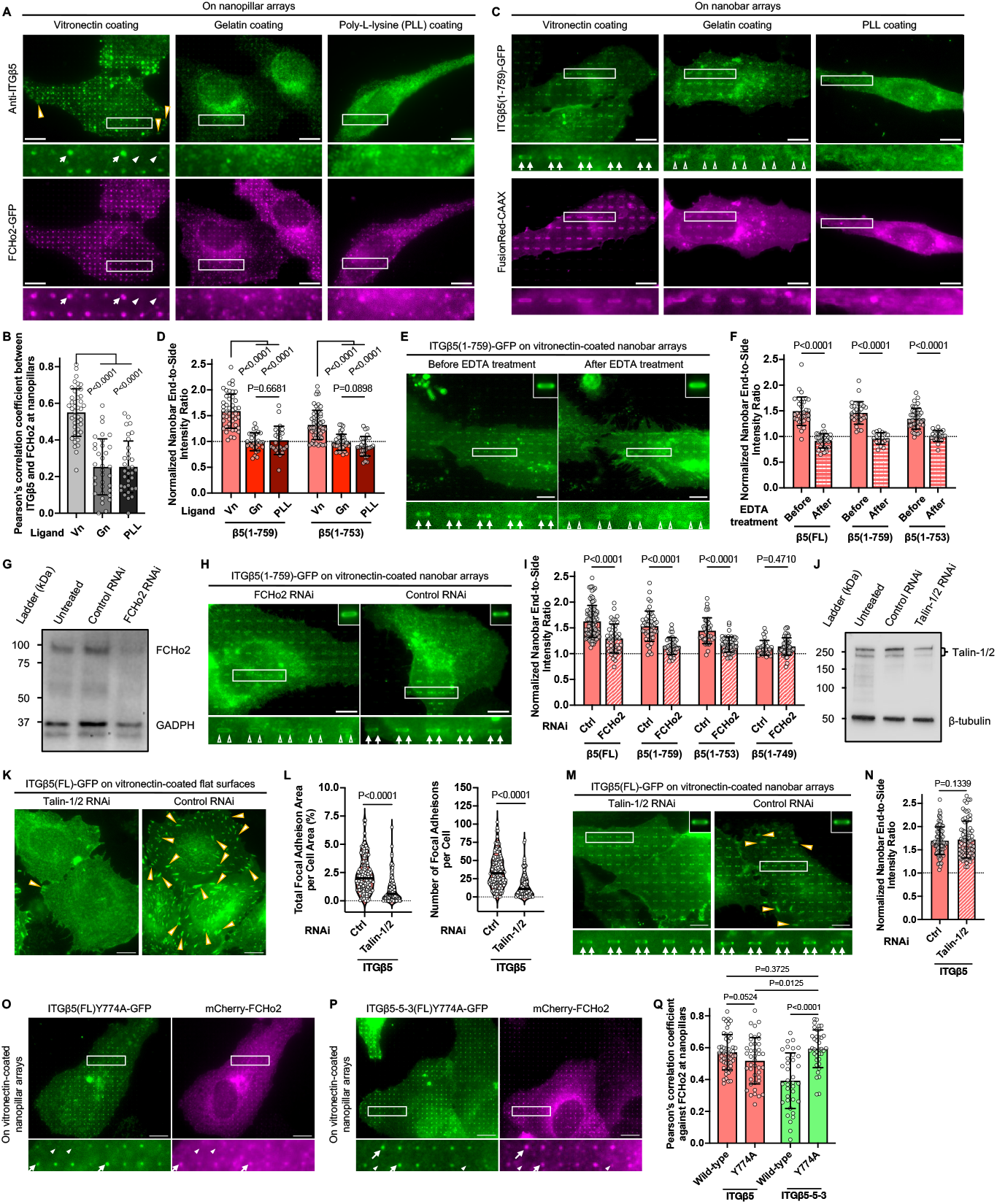
FCHo2, but not talin-1, is crucial for inside-out activation of integrin αvβ5 in curved adhesions. **(A)** Anti-ITGβ5 and FCHo2-GFP accumulate and colocalize at nanopillars when substrates are coated with vitronectin (left), an ECM ligand for integrin αvβ5, but not on substrates coated with hydrolyzed collagen (gelatin, middle), or poly-L-lysine (PLL, right), which are not the ligands for integrin αvβ5. **(B)** Quantifications of Pearson’s correlation coefficients between anti-ITGβ5 and FCHo2-GFP at nanopillars coated with different ligands. **(C)** ITGβ5(1-759)-GFP displays a clear preference for nanobar ends only when substrates are coated with vitronectin. **(D)** Quantifications of the normalized nanobar end-to-side intensity ratios of ITGβ5(1-759)-GFP and ITGβ5(1-753)-GFP on nanobar substrates coated with different ligands. Vn: vitronectin; Gn: gelatin; PLL: poly-L-lysine. **(E)** After 5-min incubation, EDTA dramatically reduced the accumulations of ITGβ5(1-759)-GFP at nanobar ends. **(F)** Quantifications of the normalized nanobar end-to-side intensity ratio of ITGβ5(FL)-GFP, ITGβ5(1-759)-GFP, and ITGβ5(1-753)-GFP before and after EDTA treatment. **(G)** Western blots confirm the efficient knockdown of FCHo2 by shRNA. **(H)** Preferential accumulations of ITGβ5(1-759)-GFP at nanobar ends are eliminated upon knockdown of FCHo2, but not by scramble (control, Ctrl) shRNA treatment. **(I)** Quantifications of the normalized nanobar end-to-side intensity ratio of four ITGβ5-GFP truncation variants upon treatments of FCHo2 shRNA or scramble shRNA. **(J)** Western blots confirm the double knockdown of talin-1 and talin-2 by shRNA. **(K)** ITGβ5-marked focal adhesions are drastically diminished upon double knockdown of talin-1 and talin-2 (left images), but retained upon treatment with control shRNA (right images). **(L)** Quantifications of the focal adhesion area percentage (left) and the number of focal adhesions per cell (right) of ITGβ5-GFP upon treatments of talin-1/2 or scramble shRNA. **(M)** Curved adhesions, illustrated by the ITGβ5-GFP enrichments at nanobar ends, remain intact upon double knockdown of talin-1 and talin-2. **(N)** Quantifications of the normalized nanobar end-to-side intensity ratio of ITGβ5-GFP upon treatments of talin-1/2 shRNA or scramble shRNA. **(O-P)** On nanopillar arrays, both ITGβ5(Y774A)-GFP (O) and ITGβ5-5-3(Y774A)-GFP (P) are spatially correlated with mCherry-FCHo2 at nanopillars. **(Q)** Quantifications of Pearson’s correlation coefficients between mCherry-FCHo2 and ITGβ5-GFP, ITGβ5-5-3-GFP, or their Y774A mutants, at nanopillars. Scale bar: 10 µm for all the cell images. White arrows indicate enrichments at nanobar ends, white empty arrowheads indicate no preferential enrichment at nanobar ends, while yellow arrowheads indicate focal adhesions. In (A) and (O)-(P), white arrows indicate high-intensity correlations, while white triangles indicate low-intensity correlations at nanopillars. In (D), (F), (I), and (N), the dashed line indicates a value of 1. Welch’s t tests (unpaired, two-tailed, not assuming equal variance) are applied for statistical analyses of curvature-related measurements, while Kolmogorov-Smirnov test was used for statistical analyses of focal adhesions. Error bars represent standard deviations. Each data point represents one cell.

**Figure 6.**
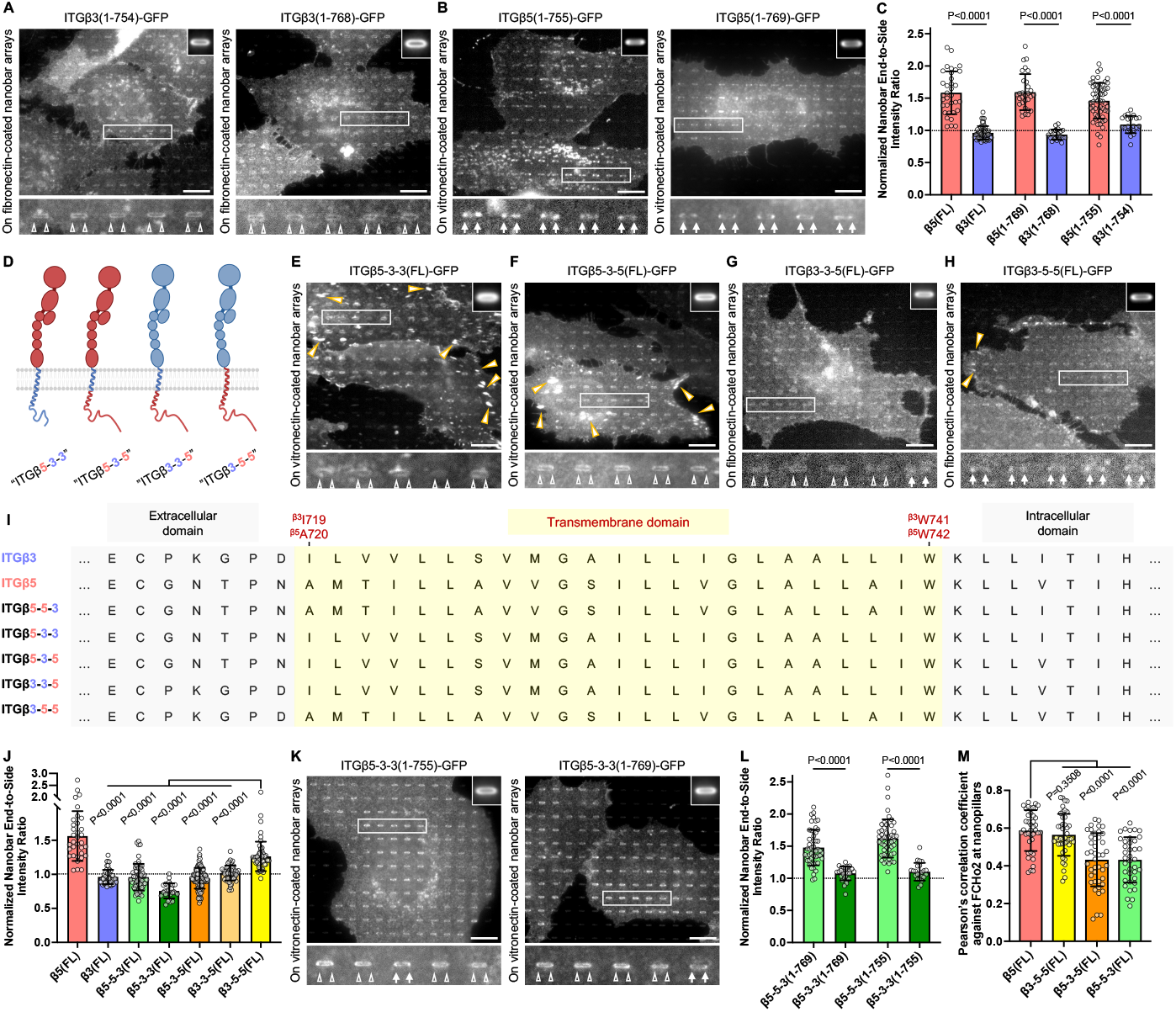
The transmembrane domain of integrin β5, but not that of integrin β3, is compatible with curvature sensitivity. **(A-B)** ITGβ3 truncation variants, ITGβ3(1-754)-GFP (A, left) and ITGβ3(1-768)-GFP (A, right) both display no curvature preference, in strong contrast to their matching ITGβ5 truncation variants, ITGβ5(1-755)-GFP (B, left) and ITGβ5(1-769)-GFP (B, right), respectively, both of which preferentially accumulate at nanobar ends (white arrows). **(C)** Quantifications of the normalized nanobar end-to-side intensity ratios of ITGβ3-GFP truncation variants and their matching ITGβ5-GFP truncation variants. **(D)** Cartoon illustration of four new integrin β5/β3 chimeric variants, including ITGβ5-3-3, ITGβ5-3-5, ITGβ3-3-5, and ITGβ3-5-5. **(E-H)** Chimeric ITGβ5-3-3-GFP (E) and ITGβ5-3-5-GFP (F) form abundant focal adhesions between nanobars but do not accumulate at nanobar ends. ITGβ3-3-5-GFP (G) forms sparse focal adhesions and does not accumulate at nanobar ends. Although ITGβ3-5-5-GFP (H) forms few focal adhesions, it displays a clear preference for nanobar ends (white arrows). **(I)** Sequence alignment of the transmembrane domains of ITGβ3, ITGβ5, and their chimeric variants. **(J)** Quantifications of the normalized nanobar end-to-side intensity ratios of seven ITGβ5/β3-GFP chimera. **(K)** ITGβ5-3-3 truncation variants, ITGβ5-3-3(1-755)-GFP (left) and ITGβ5-3-3(1-769)-GFP (right), which are devoid of the NPLY talin-binding sites, do not accumulate at nanobar ends. **(L)** Quantifications of the normalized nanobar end-to-side intensity ratios of ITGβ5-3-3-GFP truncation variants and their matching ITGβ5-5-3-GFP truncation variants. **(M)** Quantifications of Pearson’s correlation coefficients between mCherry-FCHo2 and ITGβ5-GFP, ITGβ3-5-5-GFP, ITGβ5-3-5-GFP, or ITGβ5-5-3-GFP, at nanopillars. Scale bar: 10 µm for all the cell images. White arrows indicate enrichments at nanobar ends, white empty arrowheads indicate no preferential enrichment at nanobar ends, while yellow arrowheads indicate focal adhesions. In (A)-(B), (E)-(H), and (K), the averaged nanobar images are shown on the top-right corners. In (C), (J), and (L), the dashed line indicates a value of 1. Welch’s t tests (unpaired, two-tailed, not assuming equal variance) are applied for statistical analyses of curvature-related measurements. Error bars represent standard deviations. Each data point represents one cell.

Next, we examined whether septin-2 and septin-7, two cyto-skeletal proteins implicated in integrin-mediated adhesions and known to sense membrane curvature, are involved in curved adhesions. Furthermore, FCHo2 has reportedly interacted with septins and enhanced septin bundling *in vitro*^*80*^. In agreement with previous findings, on flat substrates, septin-2 forms actin-like long filaments connected to focal adhesions, whereas septin-7 forms both long filaments and short, thick ones (**Supplementary Figure 5F-5G**)^78^. On nanobars, both septin-2 and septin-7 accumulate at nanobar ends (**Figure 3A**; Quantifications in **Figure 3B**), consistent with their known curvature preference^81–83^.

To further examine whether any of these components, particularly septin-2 and septin-7, are involved in curved adhesions, we measured their spatial correlations with ITGβ5 and FCHo2 at curved membranes. For this study, we utilized vertical nanopillar arrays (200 nm in diameter, 1 µm in height, and 2.5 µm in spacing), which induce cylindrical curvatures along the vertical heights (**Figure 3C** and **Supplementary Figure 6B**)^54,57,60^ Both nanobars and nanopillars generate well-defined membrane curvature, but they are used for different analytical purposes. Nanobars provide distinct curved ends and flat side regions, making them ideal for quantifying curvature preference (i.e. nanobar end-to-side ratios). Whereas, nanopillars are distributed at a higher density, inducing more membrane curvatures per cell for protein-protein spatial correlation analysis.

On nanopillars, ITGβ5 accumulates and shows a strong spatial correlation with FCHo2, confirming the formation of curved adhesions (**Figure 3D-3G** and **Supplementary Figure 6C-6E**). Fluorescence images and quantifications of Pearson’s correlation coefficients reveal that talin-1 enriches and exhibits strong correlations with both ITGβ5 (∼0.7) and FCHo2 (∼0.5) at nanopillars (**Figure 3D**; Quantifications in **Figure 3H-3I**). Tensin-3 and kindlin-2 display minimal enrichments at nanopillars and do not spatially correlate with ITGβ5 (∼0.1-0.3) and FCHo2 (∼0.05-0.1) (Figure 3E and Supplementary Figure 6D; Quantifications in Figure 3H-3I), resembling vinculin (**Supplementary Figure 6E**; Quantifications in **Figure 3H-3I**). Very interestingly, on nanopillar substrates but not on nanobar arrays, talin-2 shows moderate enrichments at curved membranes and spatially correlates with ITGβ5 (∼0.62) and FCHo2 (∼0.4) (**Supplementary Figure 6B**; Quantifications in **Figure 3H-3I**). The dense nanopillar arrays substantially impedes focal adhesion formation, indicating that talin-2 prefers focal adhesions but can nevertheless be recruited into curved adhesions when focal adhesions are suppressed. Additionally, overexpression of tensin-3, which triggers adhesion formation on flat surfaces, impairs the formation of curved adhesions and undermines ITGβ5-FCHo2 colocalizations at nanopillars in some cells (**Supplementary Figure 6F**; Quantifications in **Supplementary Figure 6G**). More intriguingly, despite their curvature enrichments, neither septin-2 nor septin-7 at nanopillars spatially correlates with ITGβ5 (∼-0.08) and FCHo2 (∼-0.05) (**Figure 3F-3G**; Quantifications in **Figure 3H-3I**). Therefore, while septins are curvature-sensitive and preferentially accumulate at curved membranes, they are not involved in curved adhesions. The absence of other mechanosensitive components such as tensin, kindlin, and vinculin in curved adhesions accords with our previous observations that curved adhesions bear much lower mechanical forces than focal adhesions. These results also indicate that talin-1 is the primary mechanosensitive module in curved adhesions.

The interactions of talin-1 with the MP region of integrin β tails and with the inner leaflet of the plasma membrane are essential for inside-out activation of integrins in focal adhesions (**Figure 3J**)^13,19,26,28,31^. To determine whether these interactions also play a role in curved adhesions, we generated two dominant-negative talin-1 mutants based on previous studies^28,31^: the **<u>l</u>**ipid-**<u>b</u>**inding **<u>m</u>**utant (LBM) and the K316E mutant (**Figure 3K**). The talin-1(LBM) contains substitutions of six basic residues (K272A, K274Q, and R277E in the F2 subdomain; K320A, K322A, and K324A in the F3 subdomain) that demolish talin-1 binding to anionic lipids^19,28,31,84^. Talin-1(K316E) mutant abrogates the electrostatic interaction between K316 and the membrane-proximal E753 in ITGβ5 or E752 in ITGβ3^28^. Furthermore, we also constructed two dominant-active talin-1 variants: the full-length **<u>c</u>**onstitutively **<u>a</u>**ctivated talin-1 **<u>m</u>**utant (CAM) and talin-1 head **<u>d</u>**omain variant (THD, amino acids 1-435) (**Figure 3K**). The talin-1(CAM) mutant, which harbors triple mutations (M319A, T1767L, and E1770K) to abrogate the talin-1 auto-inhibition, has reportedly shown a superior capability of ac-tivating and clustering integrins in focal adhesions^73^. The talin-1(THD) variant is considered active because it lacks the autoinhibitory rod domain, yet it is only mildly effective in activating integrins since the same rod domain is required for cytoskeletal adaptor recruitments and talin-1 dimerization^73^.

Compared with the wild-type talin-1, overexpression of the two dominant-negative variants, talin-1(LBM) and talin-1(K316E), results in fewer and smaller ITGβ5-marked focal adhesions (**Figure 3L**, yellow arrowheads; Quantifications in **Figure 3M**). Furthermore, talin-1(K316E) and talin-1(LBM) are only weakly enriched in these small focal adhesion patches, confirming the importance of the MP and lipid interactions in focal adhesion formation (**Figure 3L**). On the other hand, overexpression of the constitutively activated variant talin-1(CAM) results in significantly more and larger focal adhesions, while overexpression of talin-1(THD) leads to a comparable level of focal adhesions as the wild-type talin-1, agreeing with prior observations (**Figure 3L**, yellow arrowheads; Quantifications in **Figure 3M**)^28,73^.

To determine whether these variants are incorporated into curved adhesions, we measured their spatial correlations with ITGβ5 and FCHo2 at nanopillars. Interestingly, all the talin variants are enriched at nanopillars (**Figure 3N-3O** and **Supplementary Figure 6H-6I**). For the two dominant-negative variants, talin-1(LBM) colocalizes with ITGβ5 and FCHo2 at curved membranes to a comparable extent as the wild-type talin-1 (**Figure 3N**; Quantifications in **Figure 3P-3Q**). Whereas, talin-1(K316E) shows a lower spatial correlation with ITGβ5 but similarly colocalizes with FCHo2 at nanopillars (**Supplementary Figure 6H**; Quantifications in **Figure 3P-3Q**). Intriguingly, the two dominant-active variants, talin-1(CAM) and talin-1(THD), both show reduced colocalizations with both ITGβ5 and FCHo2 at nanopillars (**Figure 3O** and **Supplementary Figure 6I**; Quantifications in **Figure 3P-3Q**). Curved adhesions, revealed by ITGβ5 enrichments at nanopillars, are not significantly perturbed by overexpression of any of the talin-1 variants (**Figure 3R**). These results indicate that, disruption of talin-1’s interactions with lipid membranes and/or the MP region of ITGβ5 or manipulation of the talin-1 activation state, selectively modulate talin-1’s potency to activate integrins in focal adhesions, but not in curved adhesions.

### FCHo2 engages the HDRRE motif of ITGβ5 to convey curvature sensitivity

We previously unveiled that the curvature sensitivity of ITGβ5 is encoded in the cytoplasmic juxtamembrane region that binds to the µHD of FCHo2^40^. To further identify the key residues responsible for ITGβ5’s curvature sensitivity, we engineered seven ITGβ5 truncation variants inclusive of ITGβ5(1-769), ITGβ5(1-759), ITGβ5(1-757), ITGβ5(1-755), ITGβ5(1-753), ITGβ5(1-751), and ITGβ5(1-749) (**Figure 4A**). These truncation variants all lack the NPLY talin-binding site and the second NxxY kindlin-binding motif^11,14,73,85^, thus cannot participate in focal adhesions (**Supplementary Figure 7A**). Nevertheless, we found that five out of the seven truncation variants, ITGβ5(1-769), ITGβ5(1-759), ITGβ5(1-757), ITGβ5(1-755), and ITGβ5(1-753) all display clear curvature preference as reflected by their preferential accumulations at nanobar ends (**Figure 4B**, white arrows). On the other hand, ITGβ5(1-751) and ITGβ5(1-749) wrap around nanobars without an apparent enrichment at nanobar ends (**Figure 4B**, white empty triangles). Quantifications show the normalized end-to-side ratios ∼1.59 for ITGβ5(FL), ITGβ5(1-769), ITGβ5(1-759), and slightly lower ratios ∼1.40 for ITGβ5(1-757), ITGβ5(1-755), and ITGβ5(1-753), all significantly greater than that of the membrane marker CAAX. On the other hand, the ratios of β5(1-751) and β5(1-749) are similar to that of CAAX ∼1.08 (**Figure 4C**). These findings highlight that the short HDRRE sequence (amino acids 749-753) is crucial for ITGβ5’s curvature sensitivity (**Figure 4A**, highlighted by a yellow shade).

**Figure 7.**
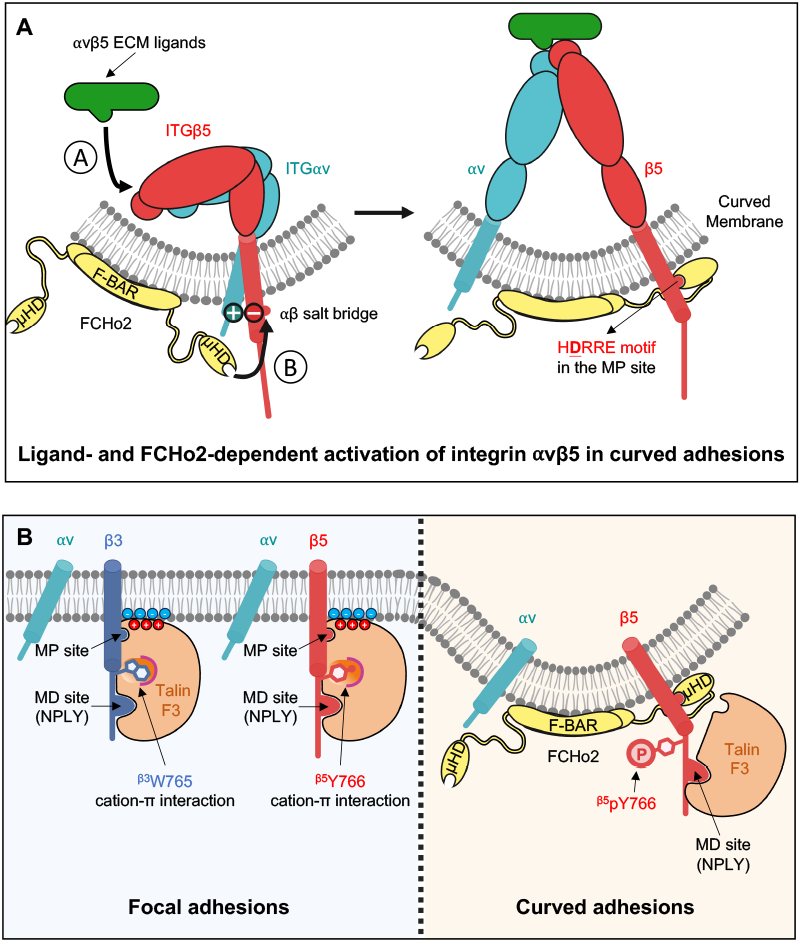
Our proposed model depicts distinctive molecular interactions and regulatory mechanisms between curved adhesions and focal adhesions. **(A)** Activation of integrin αvβ5 in curved adhesions requires both the extracellular engagement of ligands and the intracellular interaction of FCHo2 with the HDRRE motif. Upon activation, FCHo2 stabilizes liganded, active integrin αvβ5 at curved membranes. The aspartate residue (^β5^D750) of integrin β5, which forms the inhibitory salt bridge with integrin αv, is bolded and underlined. **(B)** In focal adhesions, facilitated by the favorable cation-π interaction, talin engages the integrin β tail via two contact sites: the high-affinity membrane-distal (MD) NPLY site and the low-affinity membrane-proximal (MP) site. Talin’s engagements at the MP site and with phospholipids are essential for the insideout activation of integrins in focal adhesions, which at the same time prevents FCHo2 binding to the overlapping HDRRE motif. In curved adhesions, the phosphorylation of ^β5^Y766 abolishes the cation-π interaction. This enables talin to engage only the MD site of ITGβ5 while making the HDRRE motif accessible to FCHo2 binding. Therefore, in focal adhesions, talin engages both the MD and the MP sites, and participates in both inside-out and outside-in signalings. In curved adhesions, however, talin engages the MD site and only participates in the outside-in signaling, while FCHo2 binding is essential for inside-out activation of integrin αvβ5.

The curvature preference of ITGβ5 is dependent on its interaction with a curvature-sensing protein-FCHo2^40^. To examine the role of the HDRRE motif in the ITGβ5-FCHo2 interaction, we assessed the spatial correlations between FCHo2 and different ITGβ5 truncation variants with or without the HDRRE motif, at curved membranes. We found that ITGβ5(FL), ITGβ5(1-769), ITGβ5(1-759), ITGβ5(1-757), ITGβ5(1-755), and ITGβ5(1-753), which possess the HDRRE motif, are strongly correlated with FCHo2 at curved membranes with Pearson’s correlation coefficients ∼0.5-0.58 (**Figure 4D** and **Supplementary Figure 7B**; Quantifications in **Figure 4E**). On the other hand, ITGβ5(1-751) and ITGβ5(1-749), whose HDRRE motif is partially or completely abrogated, demonstrate much lower correlations with FCHo2 at nanopillars (∼0.33), analogous to that between FCHo2 and the membrane marker CAAX (∼0.27) (**Supplementary Figure 7C**; Quantifications in **Figure 4E**). These results suggest that the HDRRE motif of ITGβ5 is crucial for FCHo2 engagement in curved adhesions.

The HDRRE motif consists of charged or polar amino acid residues. To explore the electrostatic effect, we engineered five HDRRE mutants by replacing each of the HDRRE residues with an alanine, resulting in ITGβ5(FL)H749A, ITGβ5(FL)D750A, ITGβ5(FL)R751A, ITGβ5(FL)R752A, and ITGβ5(FL)E753A. For this study, full-length ITGβ5 constructs rather than truncated versions were used in order to distinguish mutations that impair all adhesions (such as disrupting αβ pairing) from those specifically impede curved adhesion formation. Among these five mutants, ITGβ5(FL)H749A, ITGβ5(FL)R751A, and ITGβ5(FL)R752A form comparable focal adhesions as the wild-type ITGβ5 on flat surfaces, while ITGβ5(FL)E753A forms fewer focal adhesions than the wild-type (**Figure 4F**, yellow arrowheads; Quantifications in **Figure 4I**), similar to another charge inversion mutant ITGβ5(FL)E753K (**Supplementary Figure 4C**, yellow arrowheads; Quantifications in **Figure 4I**). The inhibitory effect of E753A and E753K mutations on focal adhesions is expected since the interaction between E753 in the MP region of ITGβ tail and ^talin-1^K316 is crucial for inside-out integrin activation in focal adhesions (Figure 3L-3M)^28^. Interestingly, all the five alanine mutants and the E753K mutant demonstrate drastically reduced enrichments at nanobar ends in comparison with the wild-type ITGβ5 (**Figure 4F** and **Supplementary Figure 4D**, white empty triangles), reflected by lower normalized end-to-side ratios than the wild-type ITGβ5 (∼1.09-1.23 versus ∼1.58, **Figure 4H**).

The D750 in ITGβ5 plays a crucial role in forming the inhibitory αβ salt bridge with the arginine (R995) of ITGαv subunit^86,87^. Intriguingly, ITGβ5(FL)D750A fails to form both curved adhesions and focal adhesions (**Figure 4F**, white empty triangles and yellow arrowheads; Quantifications in **Figure 4I**). It’s likely that the hydrophobic alanine side chain might perturb the membrane packing. To get further insight into this residue, we engineered ITGβ5(FL)D750R and ITGβ5(FL)D750E by replacing the negatively charged aspartate (D) with either a cationic arginine (R) or an anionic glutamate (E). The D750R mutation is expected to disrupt the inhibitory αβ salt bridge, rendering ITGβ5(FL)D750R constitutively activated. We observed that both ITGβ5(FL)D750R and ITGβ5(FL)D750E form abundant focal adhesions (Figure 4G, yellow arrowheads; Quantifications in Figure 4I). Strikingly, neither ITGβ5(FL)D750R nor ITGβ5(FL)D750E is able to form curved adhesions (**Figure 4G**, white empty triangles; Quantifications in **Figure 4H**). By measuring the spatial correlations between FCHo2 and ITGβ5 D750A/E/R mutants at nanopillars, we found that FCHo2 exhibits significantly reduced interactions with all these mutants in contrast to the wild-type ITGβ5, at nanopillars (**Supplementary Figure 7D**; Quantifications in **Figure 4J**). These results indicate that disrupting the αβ salt bridge itself is not sufficient for the formation of curved adhesions, while FCHo2 binding to the HDRRE region is crucial.

Collectively, these observations indicate that each of the five residues in the HDRRE motif is crucial for ITGβ5’s interaction with FCHo2 and its curvature preference. Several mutations, including ITGβ5(FL)H749A, ITGβ5(FL)D750E, ITGβ5(FL)D750R, ITGβ5(FL)R751A, and ITGβ5(FL)R752A, do not influence the formation of focal adhesions, but they selectively abolish the formation of curved adhesions. These results further support that the HDRRE motif is involved in different sets of molecular interactions in curved adhesion and focal adhesions.

### FCHo2, but not talin-1, is crucial for inside-out activation of integrin αvβ5 in curved adhesions

Previous studies have revealed that talin binding to integrin β intracellular domains is essential for inside-out integrin activation in focal adhesions^13,18,19,26–28^. Without talin-mediated inside-out signaling, integrins exhibit transient or low-affinity interactions with ECM ligands and cannot form focal adhesions^88^. However, our integrin β5 and β5-5-3 truncation variants lacking the NPLY talin-binding site still accumulate at curved membranes^40^. As FCHo2 engages the HDRRE motif in the membrane-proximal region of ITGβ5, a region known to be crucial for talin-mediated integrin activation, we speculate that FCHo2 binding to this region could insideout activate integrins αvβ5 in curved adhesions independent of talin.

To examine the hypothesis, we first confirmed that ITGβ5 enriched at curved membranes accommodates an active conformation that engages with extracellular ligands. To screen the effect of extracellular ligands, we coated nanobar substrates with vitronectin, gelatin, or poly-L-lysine (PLL). Among these coatings, only vitronectin serves as the extracellular ligand for integrin αvβ5^7^. ITGβ5 clusters into focal adhesions on vitronectin-coated flat substrates, yet it appears to be diffusive with cytosolic puncta on either gelatin- or PLL-coated ones (**Supplementary Figure 8A**), confirming that the formation of focal adhesions requires active integrins that engage with extracellular ligands. On nanopillar substrates, ITGβ5 strongly accumulates at nanopillars coated with vitronectin, but it displays no apparent enrichment at nanopillars with either gelatin or PLL coating (**Figure 5A**)^40^. Fluorescence recovery after photobleaching (FRAP) assay reveals that ITGβ5 at vitronectin-coated nanopillars exhibits a slow and partial recovery (<50% recovery at 5 min) (**Supplementary Figure 8B**; Quantifications in **Supplementary Figure 8C**), in contrast to the membrane marker GFP-CaaX, which recovers rapidly (∼70% recovery at 40 sec)^54^. Therefore, integrin αvβ5 stably engages with their ligands in curved adhesions. Furthermore, FCHo2 colocalizes with ITGβ5 only on vitronectin-coated nanopillar arrays (**Figure 5A**), and short-hairpin RNA (shRNA)-based RNA interferences (RNAi) of ITGβ5 significantly compromises FCHo2 accumulations at vitronectin-coated nanobar ends (Supplementary Figure 9E-9F; Quantifications in Supplementary Figure 9G). We found that FCHo2 accumulates at some gelatin- or PLL-coated nanopillars without apparent colocalizations with ITGβ5 (Figure 5A), likely related to the role of FCHo2 in clathrin-mediated endocytosis^89^. Nevertheless, FCHo2 exhibits stronger enrichments at vitronectin-coated nanobar ends over gelatin- or PLL-coated ones (**Supplementary Figure 8D**; Quantifications in **Supplementary Figure 8E**). Quantifications corroborate that FCHo2 exhibits a high spatial correlation with ITGβ5 only on vitronectin-coated nanopillar arrays (**Figure 5B**).

To examine whether integrin αvβ5 at curved membranes engages with its extracellular ligands in the absence of talin binding, we focused on ITGβ5(1-759) and ITGβ5(1-753), two representative truncation variants harboring the HDRRE motif but devoid of the NPLY talin-binding site. On the flat regions of nanobar arrays, ITGβ5(1-759) and ITGβ5(1-753) do not form focal adhesions regardless of the coatings (**Figure 5C** and **Supplementary Figure 8F**). This corroborates that talin binding is indispensable for the inside-out activation of integrin αvβ5 in focal adhesions. However, on nanobar arrays, both ITGβ5(1-759) and ITGβ5(1-753) preferentially accumulate at nanobar ends of vitronectin-coated substrates, but not of gelatin- or PLL-coated ones (**Figure 5C** and **Supplementary Figure 8F**, white arrows and white empty triangles; Quantifications in **Figure 5D**). Therefore, in the absence of talin binding, ITGβ5(1-759) and ITGβ5(1-753) can engage with their extracellular ligands selectively at curved membranes.

To further verify that integrin αvβ5 is activated in curved adhesions, we used ethylenediaminetetraacetic acid (EDTA) to inactivate integrins by sequestering divalent cations from their ligand binding pockets^90–92^. Live-cell imaging illustrates that ITGβ5(FL)-GFP, ITGβ5(1-759)-GFP, and ITGβ5(1-753)-GFP all preferentially accumulated at nanobar ends before adding EDTA (**Figure 5E** and **Supplementary Figure 8G-8I**, white arrows). After 5-min EDTA treatment, all the three variants wrap evenly around nanobars similar to the co-expressed membrane marker CAAX, with no biased distribution toward nanobar ends (**Figure 5E** and **Supplementary Figure 8G-8I**, white empty arrowheads). Quantifications of the normalized nanobar end-to-side ratios confirm that EDTA treatment abolishes the curvature enrichments of ITGβ5(FL), ITGβ5(1-759), and ITGβ5(1-753) (**Figure 5F**). Therefore, inactivation of integrins by EDTA completely eliminates the accumulation of ITGβ5 variants at curved membranes, corroborating that these integrin β5 variants accommodate an active conformation with their extracellular domains engaging with ECM ligands.

To test the hypothesis that FCHo2 is crucial for their inside-out activation at curved membranes, we exploited shRNA to knock down the endogenous FCHo2 in U2OS cells (Figure 5G). We found that FCHo2 knockdown significantly reduces the curvature enrichments of ITGβ5 variants bearing the HDRRE motif, including ITGβ5(FL)-GFP, ITGβ5(1-759)-GFP, and ITGβ5(1-753)-GFP, in comparison with the control (Ctrl; with scramble shRNA) (**Figure 5H** and **Supplementary Figure 9A-9C**, white arrows and white empty triangles). As expected, no apparent effect is observed for β5(1-749), a variant lacking the HDRRE motif that wraps around nanobars with or without FCHo2 knockdown (**Supplementary Figure 9D**, white empty arrowheads). Quantifications of the normalized nanobar end-to-side ratios confirm that, upon FCHo2 knockdown, ITGβ5(FL), ITGβ5(1-759), and ITGβ5(1-753) all exhibit significantly reduced curvature enrichments, while ITGβ5(1-749) does not accumulate at curved membranes regardless of the shRNA treatment (**Figure 5I**). Furthermore, in A549 cells that express a high level of the endogenous ITGβ5, FCHo2 knockdown largely eliminates curved adhesions at nanobar ends (**Supplementary Figure 10A**, white arrows and white empty triangles; Quantifications in **Supplementary Figure 10B**), while in the meanwhile, markedly increases viculin-positive focal adhesions, the amount of F-actin stress fibers, and overall cell size (**Supplementary Figure 10C**; Quantifications in **Supplementary Figure 10D-10G**), In contrast to FCHo2 knock-down, double knockdown of talin-1 and talin-2 leads to significantly fewer focal adhesions (**Figure 5J-5K** and **Supplementary Figure 9H**, yellow arrowheads; Quantifications in **Figure 5L**), but the enrichments of ITGβ5 at nanobar ends, namely curved adhesions, are not affected (**Figure 5M** and **Supplementary Figure 9I**, white arrows; Quantifications in **Figure 5N**). These results indicate that FCHo2, but not talin-1, plays a crucial role for inside-out integrin activation in curved adhesions, and FCHo2 knockdown enhances the formation of talin- and myosin-dependent focal adhesions.

While both talin-1 and FCHo2 interact with ITGβ5 tail and colocalize in curved adhesions, they are mutually exclusive in their interactions with the ITGβ3 intracellular domain. Unlike ITGβ5, ITGβ5-5-3 cannot form curved adhesions (**Figure 1E** and **Supplementary Figure 1B**, white empty triangles) and is not spatially correlated with FCHo2 at nanopillars (**Supplementary Figure 12C**; Quantifications in **Figure 5Q**). ITGβ5-5-3 preferentially localizes to focal adhesions where talin is enriched and FCHo2 is absent (**Figure 1F** and **Supplementary Figure 3A**). However, by abolishing talin binding, the mutant ITGβ5-5-3(Y744A) exhibits a high spatial correlation with FCHo2 at nanopillars comparable to ITGβ5 and ITGβ5-5-3(Y744A) (**Figure 4D** and 5O-5P; Quantifications in **Figure 5Q**), confirming that talin binding renders ITGβ5-5-3 curvature-insensitive by preventing FCHo2 engagement.

To examine whether FCHo2 exclusion from focal adhesions results from its competition with talin or an inability to engage flat membranes, we constructed a pan-membrane-targeting FCHo2 chimera, Lyn(SH4)-FCHo2(µHD), by replacing the F-BAR domain and IDR with the lipid-anchoring SH4 domain of Lyn (**Supplementary Figure 11A**). Neither the wild-type FCHo2 nor the Lyn(SH4)-FCHo2(µHD) chimera colocalizes with ITGβ5-marked focal adhesions on flat surfaces (**Supplementary Figure 11B-11C**). On nanobar arrays, the chimera distributes uniformly around these structures, whereas the wild-type FCHo2 selectively accumulates at curved ends (**Supplementary Figure 11D**; Quantifications in **Supplementary Figure 11E**). On nanopillar substrates, Lyn(SH4)-FCHo2(µHD) shows a substantially lower spatial correlation with ITGβ5 than the wildtype FCHo2 (**Supplementary Figure 11F-11G**; Quantifications in **Supplementary Figure 11H**). Therefore, the absence of FCHo2 from focal adhesions is not rescuable by simply providing membrane anchoring. Instead, both the F-BAR domain and µHD are required for curvature-dependent recruitment of integrin αvβ5 curved adhesions.

Taken together, our results support a model that FCHo2, instead of talin-1, is essential for inside-out activation of integrin αvβ5 in curved adhesions. Furthermore, FCHo2 binding to the HDRRE motif can be inhibited by talin-1 if the cation-π interaction forms at the pivotal location in the variants possessing W766, F766, or Y766. Abolishing this interaction, such as in the variants harboring A766, or E766, decouples talin’s engagements with the MP and MD sites, enabling FCHo2 to bind the HDRRE motif while permitting talin-1 to engage the membrane-distal NPLY motif.

### The transmembrane domain of ITGβ5, but not that of ITGβ3, is compatible with curvature sensing

In the previous sections, we employed the ITGβ5-5-3 chimera to demonstrate that talin binding to β3 intracellular domain renders it curvature insensitive. To determine whether abrogation of talin binding is sufficient to “recover” the curvature sensitivity of the wide-type ITGβ3 (ITGβ3-3-3), we engineered ITGβ3(1-768)-GFP and ITGβ3(1-754)-GFP truncation variants (**Figure 1G;** denoted by gray arrows) lacking the high-affinity talin-binding NPLY motif. Surprisingly, neither ITGβ3(1-754) nor ITGβ3(1-768) exhibits curvature preference on fibronectin-coated nanobar substrates (**Figure 6A**). In comparison, their matching ITGβ5 counterparts, ITGβ5(1-769)-GFP and ITGβ5(1-755)-GFP, both display clear curvature preferences on vitronectin-coated nanobar arrays (**Figure 6B**). The normalized end-to-side ratios of the full-length ITGβ3-GFP, ITGβ3(1-768)-GFP, and ITGβ3(1-754)-GFP are ∼1 and are much lower than their matching ITGβ5 counterparts (**Figure 6C**). These results implicate that, in addition to the intracellular talin-binding regions, the extracellular domain and/or the transmembrane domain of ITGβ3 also contribute to its insensitivity to membrane curvature.

To further investigate the distinctive curvature sensitivities between ITGβ5 and ITGβ3, we generated multiple domain-swapping chimeras in addition to ITGβ5-5-3-GFP, including ITGβ5-3-3-GFP, ITGβ5-3-5-GFP, ITGβ3-3-5-GFP, and ITGβ3-5-5-GFP (**Figure 6D**, sequence alignment in **Figure 6I**). We then compared their curvature sensitivities with the wild-type ITGβ5-GFP and ITGβ3-GFP. All 7 chimeric integrins form focal adhesions on flat areas coated with either vitronectin or fibronectin (**Figure 6E-6H**, yellow arrowheads), with ITGβ3-5-5-GFP and ITGβ3-3-5-GFP forming fewer patches than the others. Among the 7 chimeras, only ITGβ5-GFP and ITGβ3-5-5-GFP display clear curvature enrichments as confirmed by their high normalized end-to-side ratios (**Figure 6H**, white arrows; Quantifications in **Figure 6J**). A significantly lower normalized end-to-side ratio of ITGβ5-3-3 (∼0.75) is attributed to its prominent accumulations along the nanobar sidewalls (**Figure 6E**). It is particularly interesting that ITGβ5-3-5-GFP and ITGβ3-3-5-GFP present no curvature preference. The disparate behaviors between ITGβ5-GFP and ITGβ5-3-5-GFP and between ITGβ3-5-5-GFP and ITGβ3-3-5-GFP indicate that replacing the transmembrane domain of ITGβ5 with that of ITGβ3 abrogates the formation of curved adhesions. On the other hand, both ITGβ5-5-3-GFP and ITGβ5-3-3-GFP form plentiful focal adhesions, indicating that the transmembrane domains of both ITGβ5 and ITGβ3 support the formation of focal adhesions. Reduced focal adhesion formation by ITGβ3-3-5 and ITGβ3-5-5 might be attributed to the presence of the ITGβ3 extracellular domain, which exhibits a lower ligand affinity than that of ITGβ5^63^. These results further highlight that integrins in focal adhesions and curved adhesions involve distinct molecular interactions, modulated by both the intracellular and transmembrane domains.

To further examine the inhibitory effect of the ITGβ3 transmembrane domain independent of the obstructive talin-binding impact, we engineered two ITGβ5-3-3 truncation variants missing the NPLY talin-binding motif, ITGβ5-3-3(1-755) and ITGβ5-3-3(1-769), and compared their curvature sensitivities with their counterparts: ITGβ5-5-3(1-755) and ITGβ5-5-3(1-769), respectively. Both pairs differ only in their transmembrane domains. Comparison of the normalized end-to-side ratios of each pair further verify that substituting the transmembrane domain of ITGβ3 for that of ITGβ5 leads to a loss of curvature sensitivity (**Figure 6K**; Quantifications in **Figure 6L**).

Finally, we measured the spatial correlations between FCHo2 and ITGβ5, ITGβ5-3-5, ITGβ5-5-3, or ITGβ3-5-5 at nanopillars. As expected, ITGβ5 and ITGβ3-5-5, but neither ITGβ5-5-3 nor ITGβ5-3-5, spatially correlate with FCHo2 at nanopillars, indicating that the loss of curvature preference of ITGβ5-5-3 and ITGβ5-3-5 is accompanied with the loss of spatial correlation with FCHo2 (Supplementary Figure 12A-12C; Quantifications in Figure 6M). In contrast, ITGβ5, ITGβ5-3-5, and ITGβ5-5-3 form more and larger focal adhesion patches than ITGβ3-5-5 on flat substrates (**Supplementary Figure 1B** and **12D**; Quantifications in **Supplementary Figure 12E**). These results highlight that in addition to the impeding effect of ITGβ3 intracellular tail on FCHo2 binding, the transmembrane domain of ITGβ3 also attenuates FCHo2’s engagement. It is possible that the β3 trans-membrane domain is oriented in such a way that the HDRRE motif becomes inaccessible to FCHo2 binding, or that FCHo2 binding is insufficient to inside-out activate ITGβ5-3-5 to engage with extracellular ligands. Without extracellular ligand binding, FCHo2 and ITGβ5-3-5 will not be stabilized at curved membranes.

## Discussion

Talin binding to the intracellular tail of integrins has been identified as the key and committed step in inside-out integrin activation (i.e. stabilizing open, extended integrins bound to ECM ligands). Until now, talin has been the sole protein capable of performing this essential function. In this work, we show FCHo2 as an alternative to talin, capable of inside-out integrin activation selectively at curved membranes. Previous studies have established that talin’s interactions with the MP region and with anionic lipids within the cell membrane are necessary for integrin activation in focal adhesions^13,26,28^. Our studies demonstrate that disruptions of these critical interactions substantially reduce the formation of focal adhesions, but not curved adhesions, further confirming that talin-mediated inside-out activation is not essential in curved adhesions. Taken together, our data support a model, in which activation of integrin αvβ5 in curved adhesions requires both the extracellular engagement of ECM ligands and the intracellular interaction of FCHo2 with the HDRRE motif (**Figure 7A**). However, our results cannot distinguish whether extracellular ligand engagement or FCHo2 binding occurs first, a question that requires future investigations.

Our identification of a talin-independent activation mechanism of integrin αvβ5 resonates with an early study proposing a NPxY-independent ITGβ5 activation in phagocytosis^93^. In this study, Singh et. al. replaced the key tyrosine residue in NPLY with an alanine which effectively abolishes talin engagement. Interestingly, they found that this ITGβ5 mutant exhibits robust binding to microspheres coated with MFG-E8 (Milk fat globule-EGF factor 8 protein, also known as lactadherin, another ECM ligand for integrin αvβ5), indicating integrin activation. While the molecular mechanism of this phenomenon was not explored, the authors suggested the existence of a NPxY/talin-independent mechanism of integrin activation. Since this interesting study in 2007, no subsequent research has hinted at the existence of such a mechanism. On the other hand, a recent study show that the integrin αvβ5 is the most crucial integrin α/β pair for cancer cell expansion^94^. In our current work, we present compelling evidence for a talin-independent activation mechanism in curved adhesions, involving an unexpected protein FCHo2.

An intriguing aspect of curved adhesions is that only integrin αvβ5, but not the homologous αvβ3, is able to form such adhesions. On the other hand, the FCHo2-binding HDRRE motif, which is responsible for this unique capability, is not specific to integrin β5, as a similar motif is also present in integrin β3. Surprisingly, abolishing talin binding is sufficient to restore FCHo2 binding to ITGβ3 tail and its curvature preference. In our quest to unravel the underlying mechanism, we pinpointed a pivotal and highly conserved tryptophan (W) residue in the homologous integrin β isoforms -except for integrin β5, which features a tyrosine (Y) substitution at this position. Intriguingly, a single Y766W mutation renders ITGβ5(Y766W) incapable of forming curved adhesions, whereas a reverse mutation W766Y imparts ITGβ5-5-3(W766Y) chimera a new ability to form curved adhesions. Therefore, the W-to-Y substitution in β5 is crucial for its capability to form curved adhesions. Tyrosine can exist in either non-phosphorylated or phosphorylated states. Our data demonstrate that replacing the tyrosine in ITGβ5 with a phenylalanine, a non-phosphorylatable mimetic, exclusively enables the formation of focal adhesions. Whereas, substituting a phosphotyrosine mimetic glutamate for this tyrosine in ITGβ5 mostly permits the formation of curved adhesions. While our mutagenic results suggest the phosphorylation state of Y766 as a way to modulate the equilibrium between focal adhesions and curved adhesions, it has yet to be experimentally confirmed due to the lack of pY766-specific ITGβ5 antibody. Future investigations such as mass spectrometry-based proteomic studies are necessary to reveal the phosphorylation state of ^β5^Y766 in focal and curved adhesions.

We propose a model where, in focal adhesions, inside-out integrin activation is mediated by talin which engages both the MD and MP sites of integrin β tails as well as lipid membranes. While in curved adhesions, inside-out activation is mediated by FCHo2 binding to the HDRRE motif of integrin β5 with talin engaging only the MD site (**Figure 7B**). This model depicts distinct molecular interactions in focal adhesions and curved adhesions. In focal adhesions, the favorable cation-π interaction enables taline to engage both MD and MP sites while simultaneously excluding FCHo2 binding to the MP region. This agrees with our observations that FCHo2 is excluded from focal adhesions. In curved adhesions, however, talin only engages the MD site for the out-side-in signaling. These enable FCHo2 to associate with the HDRRE motif for the inside-out activation of integrin αvβ5 at curved membranes. The MD-only engagement of talin in curved adhesions likely accounts for the previous observations that talin bears much lower mechanical forces in curved adhesions (3-5 pN) than in focal adhesions (>10 pN)^40^. Since it takes > 5 pN to unravel the cryptic vinculin-binding sites in the talin rod domain^95^, the low mechanical forces in curved adhesions align with the observation that vinculin, tensin, and kindlin are absent in curved adhesions. Our model also aligns with previous observations that W-to-A and W-to-E mutations at the pivotal site reduced the formation of focal adhesions^17,96^.

Aside from the intracellular domain, the transmembrane domain of ITGβ5 is also crucial for its curvature sensitivity as well as the interaction with FCHo2. The transmembrane domain is important for both inside-out and outside-in signal transductions. An earlier study using leucine scanning revealed extensive interactions between the integrin α- and β-transmembrane domains^97^. Another study unveiled that the transmembrane domains of ITGβ3 and ITGβ1 exhibit structural differences, which modulate their associations with the ITGα subunit^98^. Indeed, there are substantial differences in the amino acid sequence of the transmembrane domains between ITGβ3 and ITGβ5. Future studies such as biophysical characterizations of protein structures, binding affinity, dynamic measurements, are warranted to further understand the regulatory mechanism behind the formation of curved adhesions.

## Acknowledgements

We thank Stanford Nanofabrication Facility (SNF) and Stanford Nano Shared Facilities (SNSF) for the nanofabrication and SEM characterization of vertical nanostructures. We thank the Cui lab members, particularly Ms. Erica Liu, for the support of this work. We thank the Bertozzi lab at Stanford Chemistry for sharing the confocal microscope. Most schematics were created with BioRender.com. This work was supported by the National Institutes of Health (R35GM141598, R01HL165491 and R01NS121934), Ono Pharma Breakthrough award (to B. C.), Stanford University Center for Molecular Analysis and Design (CMAD) fellowship (to C.-H. L.), Stanford Molecular Biophysics Training Program T32GM136568 (to C.E.L.), and Stanford Sarafan ChEM-H Chemistry/Biology Interface Program 1T32GM139791-01A1 (to L.A.V.). This paper was typeset with the bioRxiv word template by @Chrelli: www.github.com/chrelli/bioRxiv-word-template.

## Author contributions

C.-H.L. and B.C. conceptualized and designed the research; C.-H.L., performed the most experiments and analyzed all the data. C.-H.L., C.E.L., and W.Z. performed fluorescence imaging. C.-H.L., C.E.L., W.Z., and L.A.V. cloned plasmid constructs. W.Z., Y.Y., and H.Y. generated lentiviral particles for stable knockdown. C.-H.L., C.E.L. and Y.Y. performed Western blotting. C.-T.T. fabricated nanostructured substrates and took the SEM images. B.C. developed the Matlab codes for the analysis. C.-H.L. and B.C. wrote the paper. All the authors discussed the results and commented on the manuscript.

C.E.L. and W.Z. contributed equally to this work.

## Competing interest statement

All authors declare no competing interests.

## Materials and Methods

### Nanostructure fabrication and characterization

Both 200-nm nanobar and nanopillar arrays were fabricated via the two-step etching, including optical photolithography followed by reactive ion etching (RIE), as previously reported^55^. Briefly, quartz wafers were cleaned with Spin Rinse Dryer (SRD), baked, and applied with hexamethyldisilazane (HMDS) to promote photoresist adhesion. The wafers were then coated with a layer of 1-µm-thick photoresist (Shipley 3612) prior to exposure to the desired pattern of UV using Heidelberg (MLA150). The post-exposure wafers were immediately developed with the MF-26A developer (Transene). The AJA e-beam evaporator was subsequently employed to deposit a 120-nm-thick layer of chromium mask on the patterned wafers. After Cr mask deposition, photoresists were immediately lifted off with acetone and isopropanol, and the quartz wafers were etched anisotropically by RIE (Plasma-Therm Versaline LL ICP Dielectric Etcher, PT-Ox) with a mixture of C_4_F_8_, H_2_, and Ar. The quartz wafers were then incubated in 20:1 Buffered oxide etch (BOE) to isotropically shrink nanostructures to the desired dimensions after the removal of Cr mask with a chromium etchant 1020 (Transene). Nanostructured quartz wafers were then cut into several small chips for cell-based experiments. The shape and dimension of the nanostructures were measured by scanning electron microscopy (FEI Magellan 400 XHR). Detailed dimensions of the nanostructures are described in the main text and figure captions.

### Plasmid construction

Wild-type ITGβ5-EGFP (simply noted as ITGβ5-”GFP”) and wild-type ITGβ3-EGFP (noted as ITGβ3-”GFP”) were cloned as previously described^40^. Briefly, the DNA fragment encoding the human integrin β5 was amplified from the pCX-EGFP β5 integrin receptor (a kind gift from Raymond Birge, Addgene, #14996) using polymerase chain reactions (PCR). The DNA fragment encoding the human integrin β3 was amplified from the complementary DNA (cDNA) of U2OS cells. The fragments were then integrated into the pEGFP-N1 vector (Clontech) through overlap-extension PCR to obtain C-terminally tagged ITGβ5-GFP and ITGβ3-GFP, respectively. All the C-terminally integrin-GFP variants used in this work share a common flexible linker (QSTVPRARDPPVAT) which connects the integrin moiety with GFP.

EGFP-talin-1 (#14996), EGFP-talin-2 (#14996), septin-2-sfGFP (#14996), sfGFP-septin-7 (#14996), and mCherry-tensin-3 (#14996) plasmids were purchased from Addgene. The mCherry-talin-1 plasmid was a kind gift from Michael Davidson (Addgene, #55139), while mCherry-FCHo2 construct was a kind gift from Christien Merrifield (Addgene, #27686). mCherry-CAAX and FusionRed-CAAX construct were generated by inserting the DNA fragment encoding the CAAX motif of K-Ras protein (GKKKKKKSKTKCVIM) into the pFusionRed-N vector (Evrogen, #FP412) or the pmCherry-C1 vector (Clontech), respectively.

FCHo2-EGFP, Lyn(SH4)-FCHo2(µHD), and most chimeric ITGβ5/β3-GFP variants were generated via Gibson assembly. Briefly, the linear vector and the fragment were amplified by PCR. Subsequently, the fragment was inserted into the vector and then cyclized in the Gibson Assembly mixture (New England Biolabs, #E2611) composed of 2 U/µL Taq DNA Ligase, 0.025 U/µL Q5 High Fidelity Polymerase, 0.002 U/µL T5 exonuclease, and 0.05 U/µL DpnI at 50 °C for 30 min. All truncation variants and mutants of ITGβ5-GFP, ITGβ3-GFP, chimeric ITGβ5/β3-GFP, or mCherry-talin-1 were generated via site-directed mutagenesis with the aid of T4 DNA ligase (New England Biolabs, #M0202). Prior to PCR amplification of fragments for T4 ligation reactions, all primers were subject to 5’-phosphorylation by T4 polynucleotide kinase (PNK, New England Biolabs, #M0201) at 37 °C for 30 min. T4 ligation reactions were performed at 37 °C for 1 hour up to overnight.

All shRNA (small-hairpin RNA) used in this work were cloned using the protocol from Addgene “pLKO.1 Protocol”. Briefly, the fragments coding for either scramble shRNA, ITGβ5-shRNA, FCHo2-shRNA, or talin-1/2 shRNA were incorporated into the pLKO.1 vector linearized by AgeI and EcoRI. The gene encoding either BFP (for scramble, FCHo2 shRNA, and talin-1/2 shRNA) or mCherry (for ITGβ5 shRNA) was included in the vector to fluorescently mark the cells that were transiently or stably transfected. Sequence and proposed knockdown efficiency of shRNA are listed in the Supplementary Table 1.

### Substrate functionalization

Substrates were functionalized via a multilayer coating. Nanostructured chips were cleaned in a piranha solution for 2-24 hrs then by air plasma for 30-60 min. Cleaned nanostructured chips were placed into a 24-well plate, then coated with 0.2 mg/mL poly-L-lysine (PLL, Sigma-Aldrich, #P5899) for 20-30 min, 0.5% glutaraldehyde (Sigma-Aldrich, #354400) for 20 min, and 25 µg/mL vitronectin (Peprotech, #AF-140-09), fibronectin (Sigma-Aldrich, #F1141) or gelatin (Sigma-Aldrich, #G9391) for 30 min. Subsequently, the chips were incubated with 1X Dulbecco’s Modified Eagle Medium (DMEM, Gibco, #11965092) with 10% fetal bovine serum (FBS, Cytiva, #SH30071) for 30 min to quench free aldehydes. To effectively activate integrin β variants, we used vitronectin as the ECM ligand for the integrin β variants possessing the β5 extracellular domain (i.e. ITGβ5, ITGβ5-5-3, ITGβ5-3-3, ITGβ5-3-5, and their variants), and fibronectin for those harboring the β3 extracellular domain (i.e. ITGβ3 and its variants, ITGβ3-5-5, and ITGβ3-3-5).

### Plasmid transfection and cell culture

We mainly took advantage of electroporation for transient protein expressions. To harvest cells for transfection, confluent U2OS cells (ATCC, HTB-96) were first trypsinized for 5–15 min at 37 °C followed by centrifugation. The pellets were resuspended with 1X DMEM (with 10% FBS but no antibiotics) or 1X PBS. Second centrifugation was applied to remove residual trypsin and EDTA. A half 6-well well of confluent U2OS cells was used for one electroporation reaction. For all ITGβ5- and ITGβ3-GFP variants, ∼500 ng of plasmid DNA was mixed in a solution composed of 2 µL of electroporation buffer I (360 mM adenosine 5′-triphosphate and 600 mM magnesium chloride) and 100 µL of electroporation buffer II (88 mM monobasic potassium phosphate and 14 mM sodium bicarbonate at pH = 7.4). If desired, ∼1.5 µg of plasmid DNA encoding either scramble, ITGβ5, or FCHo2 shRNAs were included in the mixture. Cell pellets were gently resuspended with the electroporation mixture and subsequently transferred into a 0.2-cm electroporation cuvette (Thermo Fisher Scientific, #FB102). The cell-DNA mixture was then electroporated with a U2OS-specific program using Amaxa Nucleofector II (Lonza). To complete the electroporation, the mixture was immediately added with 800-1000 µL of pre-warmed (37 °C) 1X DMEM (with 10% FBS but no antibiotics) and incubated for 10-15 min. Cells were then harvested by spinning and resuspended with 1X DMEM with 10% FBS, 100 U/mL penicillin and 100 mg/mL streptomycin (Gibco, #15140122). The transfected cells were cultured in a 6-well or 12-well plate, and maintained in a standard incubator at 37 °C with 5% CO_2_ for 72 hrs.

After 72-hr incubation, ITGβ-GFP-transfected U2OS cells were detached from well plates with a pre-warmed 5 mM EDTA in 1X PBS at 37 °C for 15-20 min. Cells were then harvested via spinning. If needed, 72-hr-cultured U2OS cells expressing a given ITGβ-GFP variant were further co-transfected with ∼500 ng of the constructs for mCherry-FCHo2, mCherry-talin-1, or their variants, via electroporation. After resuspending cell pellets with 1X DMEM supplemented with 10% FBS and antibiotics, desired amounts of U2OS cells were then plated on the functionalized nanostructured or flat substrates and maintained in with 1X DMEM (with 10% FBS and antibiotics). The cultures were incubated in a standard incubator for 24 hrs at 37 °C with 5% CO_2_. Upon 96-hr incubation (72-hr post-transfection incubation plus 24-hr plating/culture on substrates), the majority of ITGβ-GFP signals were localized to the plasma membrane and adhesion architectures, with a small residual amount trapped in the endoplasmic reticulum (ER) or perinuclear regions. Cells exhibiting strong ER-localized ITGβ-GFP signals were excluded from all analyses.

For overexpression of GFP-talin-1, GFP-talin-2, mCherry-tensin-3, septin-2-GFP, GFP-septin-7, or FusionRed-CAAX, 300-400 ng of plasmids were used for electroporation. If integrin β-GFP overexpression was omitted (for instance, endogenous ITGβ5 immunostaining experiments), 24-hr post-transfection culture on substrates was sufficient.

### Cell membrane visualization

To visualize the plasma membrane in U2OS cells, we mostly performed transient expression of the fluorescent CAAX (FusionRed-CAAX). 72-hr-cultured U2OS cells transfected with a given ITGβ-GFP variant were further co-transfected with ∼300 ng of the FusionRed-CAAX plasmid via electroporation. For three-channel imaging, we transiently transfected U2OS cells with an extracellular SNAP-tag to label the plasma membrane with Alexa Fluor 647. Briefly, cells were transfected with SNAP-pDisplay and then incubated in a pre-warmed 1X DMEM medium supplemented with 1 µM O^6^-benzylguanine (BG)-coupled Alexa Fluor 647 (NEB, #806 S9136S) at 37 °C with 5% CO_2_. After 15-min incubation, the samples were washed 5 times with a pre-warmed 1X DMEM medium before fixation and imaging.

### Cell fixation and immunostaining

To probe the endogenous ITGβ5, overnight-seeded U2OS cells (on either nanostructured substrates or flat surfaces) were fixed in 4% paraformaldehyde (PFA) (Thermo Fisher Scientific, #28908) in 1X PBS at room temperature for 15 min. To preserve focal adhesion architectures marked by cytoskeletal adaptors (i.e. talin-1, talin-2, tensin-3, kindlin, and vinculin), cells were instead fixed in 4% PFA in 1X PHEM buffer (60 mM PIPES, 25 mM HEPES, 10 mM EGTA, 2 mM MgCl_2_, pH=7.0) at room temperature for 5-10 min. After three washes with 1X PBS, cells were then permeabilized and blocked with 0.1% Triton X-100 and 1% bovine serum albumin (BSA) (Sigma-Aldrich, #A9418) in 1X PBS (staining buffer) at room temperature for 1 hr. Cells were then incubated with rabbit anti-ITGβ5 (Cell Signaling, clone D24A5, #3629S), mouse anti-kindlin-2 antibody (Antibodies.com, clone 3A3, #A277536), or mouse anti-vinculin antibody (Sigma-Aldrich, clone hVIN-1, #V9131) at 1:500 dilution in the staining buffer for 0.5-1 hr at room temperature. After three washes with 1X PBS, cells were then stained with secondary antibodies (goat anti-rabbit IgG Alexa Fluor 647 (Invitrogen, #A-32733), goat anti-mouse IgG Alexa Fluor 488 (Invitrogen, #A-32723), or goat anti-mouse IgG Alexa Fluor 594 (Invitrogen, #A-11032)) at 1:500 dilution in the staining buffer for 0.5-1 hr at room temperature in the dark. Samples were washed with 1X PBS three times prior to fluorescence imaging.

For A549 cell experiments, blank (untreated, wild-type) cells were mixed with FCHo2-depleted cells, then subsequently plated onto vitronectin-coated nanobar arrays or flat substrates. 4 hours post plating and prior to fluorescence imaging, cells were fixed, permeabilized, and stained with phalloidin (Invitrogen, Alexa Fluor 647-labeled, #A22287), rabbit anti-ITGβ5, and mouse anti-vinculin.

The antibodies used in this work are listed in the **Supplementary Table 2**.

### Mn^2+^ treatment experiments

Overnight-seeded ITGβ-GFP-expressing U2OS cells were incubated with 0.5 mM Mn(NO3)_2_ (Supelco, #1.19789) in a pre-warmed DMEM supplemented with 10% FBS and antibiotics for 20 min at 37 °C. Upon completion of the treatment, cells were washed and fixed prior to imaging.

### EDTA treatment experiments

For EDTA treatment experiments, live cell assays were performed. Prior to EDTA addition, overnight-seeded ITGβ-GFP-expressing U2OS cells were first imaged to record “before” signals. Cells were then treated with 5 mM EDTA (Invitrogen, #15575020) in a pre-warmed DMEM supplemented with 10% FBS and antibiotics for 4-5 min at 37 °C, and the samples were imaged at 37 °C in a microscopic chamber supplemented with 5% CO_2_ to record “after” signals.

### Lentiviral particle generation for stable shRNA knockdown

We followed and modified the protocol for lentiviral particle generation from Addgene “pLKO.1 Protocol”. Briefly, HEK293T cells were cultured in 1X DMEM supplemented with 10% FBS and antibiotics in six-well plates to reach ∼70–80% confluency. Prior to transient transfection, the growth medium was replaced with pre-warmed opti-MEM. Subsequently, HEK293T cells were transiently transfected with 0.8 µg psPAX2 plasmid, 0.7 µg pMD2.G plasmid, and 1.5 µg transfer plasmid encoding pLKO.1-shRNA-BFP constructs using Lipofectamine 2000. The transfection mixture was replaced with fresh 1X DMEM (with 10% FBS, 1 mM sodium pyruvate (11360; Gibco), and antibiotics) 6 hrs post transfection. The virus-containing supernatants were collected and filtered through 0.45-µm polyvinylidene difluoride filters (Millipore) 24-48 hrs after transfection. Target cells were transduced with lentivirus and incubated for at least 48-72 hrs prior to fluorescence imaging experiments or Western blot analysis.

### Western blot analysis for shRNA knockdown

U2OS cells were transduced with lentivirus harboring shRNA of interest and incubated at 37 °C with 5% CO_2_ for 2-3 days. 24 hrs post transduction, virus-containing growth medium was removed and replaced with fresh 1X DMEM (with 10% FBS and antibiotics), allowing cells to grow for 1-2 more days (2-3 days in total). Prior to lysis, cells were detached with EDTA, harvested by spinning, and rinsed with ice-cold 1X PBS. The cells pellets were resuspended and lysed in 1X RIPA buffer (25 mM Tris-HCl, 150 mM NaCl, 1% Triton X-100, 1% sodium deoxycholate, and 0.1% sodium dodecyl sulfate) supplemented with protease and phosphatase inhibitor cocktails (Roche, #04693159001 and #04906837001) for 1 hr at 4 °C. For probing the endogenous ITGβ5, the samples were further boiled for 10 min at 95°C. The clarified lysates were mixed with 4X Laemmli sample buffer (Bio-Rad, #1610747) and β-mercaptoethanol, and boiled for 5 min at 95°C. The samples were subsequently resolved by SDS-PAGE using Mini-Protean vertical electro-phoresis system (Bio-Rad, #1658026FC) for 60-70 min at 100-110 V. After SDS-PAGE, the samples were then transferred onto a nitrocellulose membrane using a Trans-Blot Turbo Transfer system (Bio-Rad, #1704150) following manufacturer’s instructions. The membranes were blocked with 3% BSA in 1X TBST buffer for 30 min at room temperature and then incubated with primary antibodies (rabbit anti-ITGβ5 (Cell Signaling, clone D24A5, #3629S), rabbit anti-FCHo2 (Novus Biologicals, Polyclonal, #NBP2-32694), or rabbit anti-GAPDH (Cell Signaling, clone 14C10, #2118) overnight at 4°C. After 1-hr incubation with HRP-conjugated secondary antibodies at room temperature, the protein bands were visualized through chemiluminescent signals by Azure Imaging Systems (Azure Biosystem).

### Fluorescence imaging

Fluorescence imaging was performed using an epi-fluorescence microscope (Leica DMI 8000) controlled by the Leica LAS X Leica LED8 system. Nanostructured chips were flipped and placed on a glass-bottom petri dish when imaging cells^55^. The samples were imaged with a 63X oil immersion objective (N.A.=1.4) and acquired using the K8 Scientific CMOS microscope camera.

### Fluorescence recovery after photobleaching (FRAP) assay

FRAP was performed using a confocal microscope (Nikon A1plus) controlled by NIS-Elements AR. The ITGβ5-GFP signals were imaged with a 488 nm laser using a 63X oil immersion objective (N.A.=1.4). 1-µm-diameter circular ROIs at nanopillars were bleached for 2 sec using the 488-nm at ∼10% power. Pre-bleaching (-5 to -10 frames) and post-bleaching time courses were acquired at 1 sec/frame for around 5 minutes.

### Quantification of normalized nanobar end-to-side intensity ratios

The details and flowchart for the analysis are described in our previous work^40,56^. Briefly, fluorescence images were processed and analyzed using MATLAB (Version 2021a) and Fiji (ImageJ2, Version 2.3.0). We use a custom-written MATLAB code to generate a matrix of masks to include each nanobar covered by cells in the bright-field channel. These masks were then applied to the fluorescence channels to create 40^*^40-pixel averaged images of proteins of interest (POIs) on nanobars. Background subtraction using the rolling ball algorithm with a radius of 1000 pixels was applied to the average images before quantifying nanobar end-to-side ratios. We used a 3^*^3-pixel square to cover the regions of interest (ROIs) on a nanobar, including nanobar ends and flat sidewalls (**Supplementary Figure 1C**). The nanobar end-to-side intensity ratios from multiple nanobars in an individual cell were averaged and are reported as a single data point. Based on the integrity of nanostructures and the location of a cell adhered to nanostructured substrates, a single cell may cover a wide range of numbers of nanobars from ∼30 nanobars/cell to ∼300 nanobars/cell.

### Quantification of spatial correlations at nanopillars

The details for the analysis are described in our previous work^40^. To quantify the spatial correlation between two POIs at nanopillar locations, the individual fluorescence images of POIs, rather than the averaged ones, were extracted using the abovementioned custom-written MATLAB code. After background subtraction using the rolling ball algorithm with a radius of 1000 pixels, the nanopillar ROIs were selected by a 7-pixel-diameter circle and the mean fluorescence intensities of POIs at individual nanopillars were measured using Fiji. The degree of spatial correlation between two arrays of POIs’ intensities at individual nanopillars in a cell, reflected by the Pearson’s correlation coefficients, was calculated using Prism 9 (GraphPad Software). The Pearson’s correlation coefficients measured from each cell are reported as a single data point. Based on the integrity of nanostructures and the location of a cell adhered to nanostructured substrates, a single cell may cover a wide range of numbers of nanopillars from ∼60 nanopillars/cell to ∼700 nanopillars/cell.

### Quantification of normalized nanopillar-to-surrounding intensity ratios

The details and flowchart for the analysis are described in our previous work^40,56^. The analysis pipeline resembles that for nanobar end-to-side intensity ratio quantifications. The 20^*^20-pixel averaged images of proteins of interest at nanopillars were background-subtracted. A 7-pixel-diameter circle was then employed to cover the nanopillar ROIs, with the rest of areas serving as the surrounding ROIs (**Supplementary Figure 6B**). The nanopillar-to-surrounding intensity from multiple nanopillars in an individual cell were averaged and reported as a single data point.

### Quantification of focal adhesions on flat substrates

We adapted and modified the analysis method for quantifying focal adhesions from a previous work^99^. All the processes were performed using Fiji. Briefly, the ROIs were first selected using the wand tool. Fluorescence images were background-subtracted by the rolling ball algorithm (radius = 25 pixels, sliding paraboloid). Subsequently, we ran the CLAHE plug-in to enhance the local contrast of the images (block size = 9, histogram bins = 128, maximum slope = 6, no mask and no fast), followed by mathematical exponential (EXP) to further reduce backgrounds. The brightness and contrast of the images were then adjusted automatically, and thresholding was performed using the Huang method. Eventually, we quantified the focal area percentage (Total focal adhesion area per cell area) and the number of focal adhesions per cell through the ANALYZE PARTICLES function (size = 0.5-25 µm^2^, circularity = 0.00-0.99, no Exclude on edges).

### Statistics and reproducibility

Parametric, Welch’s t-tests (unpaired, two-tailed, not assuming equal variance) was employed to evaluate the statistical significance for the quantifications of curvature-related measurements (i.e. normalized nanobar end-to-side intensity ratios, Pearson’s correlation coefficients at nanopillars, etc.), while nonparametric, Kolmogorov-Smirnov test was used for assessing the statistical difference for the quantifications of focal adhesions (i.e. total focal adhesion area per cell area and number of focal adhesions per cell). All data were presented as mean ± SD. All statistical analyses were performed using Prism 9. All experiments were repeated at least 2-3 times to ensure reproducibility. All measurements were taken from distinct samples; no sample was measured repeatedly.

## Notes

### Competing Interest Statement

The authors have declared no competing interest.

### Summary of Updates

We have modified the authors' affiliation.

