## Supplementary information for "FCHo2, instead of talin, enables inside-out activation of integrin αvβ5 in curved adhesions"

bioRxiv

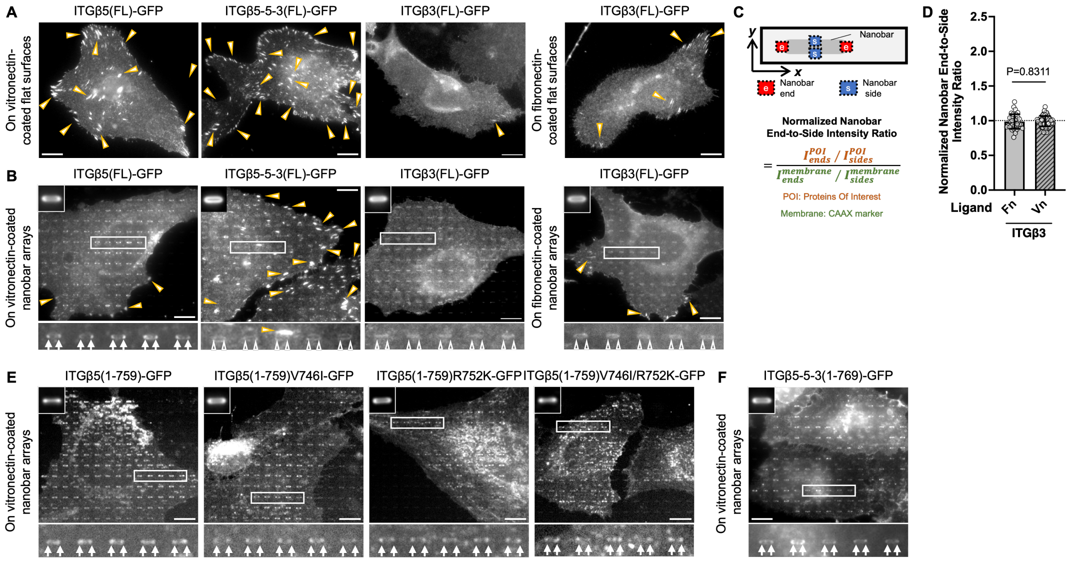


**Supplementary Figure 1. Talin binding prevents the intracellular domain of ITGβ3, but not that of ITGβ5, from responding to membrane curvature (Related to Figure 1). (A-B)** Representative images of ITGβ5-GFP, ITGβ5-5-3-GFP, and ITGβ3-GFP on flat substrates (A) or nanobar arrays (B). All the three integrins form focal adhesions on flat areas, but only ITGβ5-GFP demonstrates a clear curvature preference and accumulates in curved adhesions formed at nanobar ends. ITGβ3-GFP does not show an apparent curvature sensitivity on nanobar substrates, regardless of extracellular ligand coatings. **(C)** Schematic illustrations depicting how normalized nanobar end-to-side intensity ratios are quantified. **(D)** Quantifications of the normalized nanobar end-to-side intensity ratios of ITGβ3-GFP on fibronectin- and vitronectin-coated nanobar arrays. Fn: fibronectin; Vn: vitronectin. The dashed line indicates a value of 1. **(E)** ITGβ5(1-759)GFP, ITGβ5(1-759)V746I-GFP, ITGβ5(1-759)R752K-GFP, and the double mutant ITGβ5(1-759)V746I/R752K-GFP all clearly display preferential accumulations at nanobar ends. **(F)** ITGβ5-5-3(1-769)-GFP preferentially accumulates at nanobar ends. Scale bar: 10 μm for all the cell images. White arrows indicate enrichments at nanobar ends, white empty arrowheads indicate no preferential enrichment at nanobar ends, while yellow arrowheads indicate focal adhesions. In (B) and (E)-(F), the averaged nanobar images are shown on the top-left corners. Welch's t tests (unpaired, two-tailed, not assuming equal variance) are applied for statistical analyses of curvature-related measurements (i.e. normalized nanobar end-to-side intensity ratios). Error bars represent standard deviations. Each data point represents one cell.

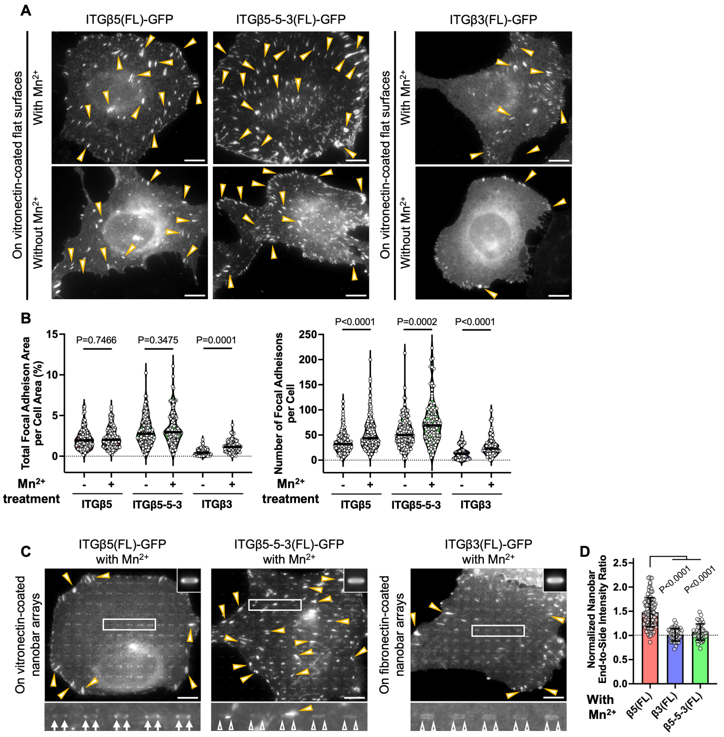


**Supplementary Figure 2. $Mn^{2+}$ treatments do not alter the curvature sensitivities of ITGβ5, ITGβ5-5-3, and ITGβ3 (Related to Figure 1). (A)** Representative images of ITGβ5-GFP, ITGβ5-5-3-GFP, and ITGβ3-GFP on flat substrates, in the presence (top panel) or absence of $Mn^{2+}$ (bottom panel). **(B)** Quantifications of the focal adhesion area percentage (left) and the number of focal adhesions per cell (right) of the three β integrin-GFP, with or without $Mn^{2+}$ treatment. **(C)** Representative images of ITGβ5-GFP, ITGβ5-5-3-GFP, and ITGβ3-GFP on nanobar arrays in the presence of $Mn^{2+}$. The averaged nanobar images are shown on the top-right corners. **(D)** Quantifications of the normalized nanobar end-to-side intensity ratios of the three β integrin-GFP in the presence of $Mn^{2+}$. The dashed line indicates a value of 1. Scale bar: 10 μm for all the cell images. White arrows indicate enrichments at nanobar ends, white empty arrowheads indicate no preferential enrichment at nanobar ends, while yellow arrowheads indicate focal adhesions. Welch's t tests (unpaired, two-tailed, not assuming equal variance) are applied for statistical analyses of curvature-related measurements, while Kolmogorov-Smirnov test was used for statistical analyses of focal adhesions. Error bars represent standard deviations. Each data point represents one cell.

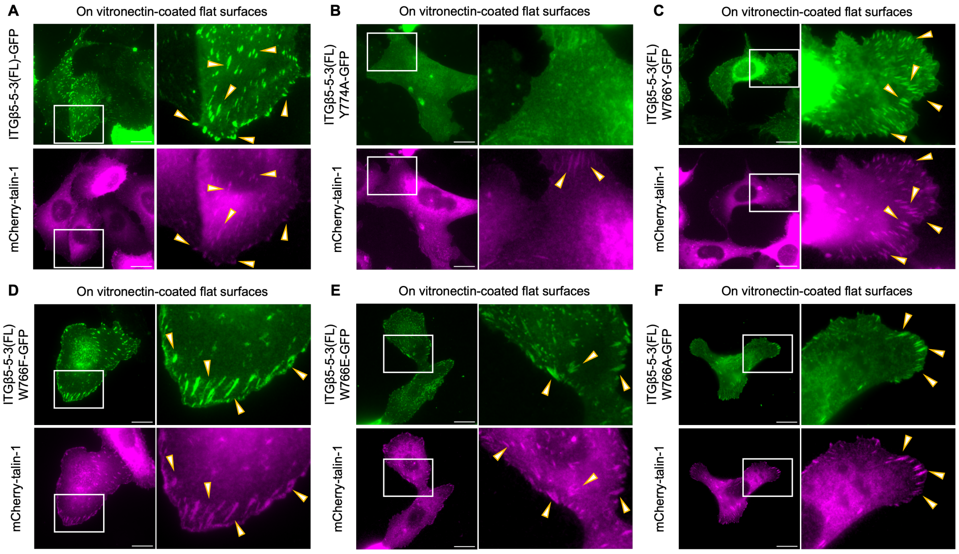


**Supplementary Figure 3. Co-expression of mCherry-talin-1 and ITGβ5-5-3-GFP mutants on vitronectin-coated flat substrates (Related to Figure 2).** **(A-F)** Representative images of the wild-type ITGβ5-5-3-GFP (A), ITGβ5-5-3(W766Y)-GFP (C), ITGβ5-5-3(W766F)-GFP (D), ITGβ5-5-3(W766E)-GFP (E), and ITGβ5-5-3(W766A)-GFP (F), which all colocalize with mCherry-talin-1 in focal adhesion patches on flat substrates (yellow arrowheads). However, ITGβ5-5-3(Y774A)-GFP (B) is absent from mCherry-talin1-marked focal adhesions. In ITGβ5-5-3(Y774A)-GFP-expressing cells, the amount and size of focal adhesions marked by mCherry-talin 1 are also significantly reduced. Scale bar: 20 μm for all the images. Yellow arrowheads indicate focal adhesions.

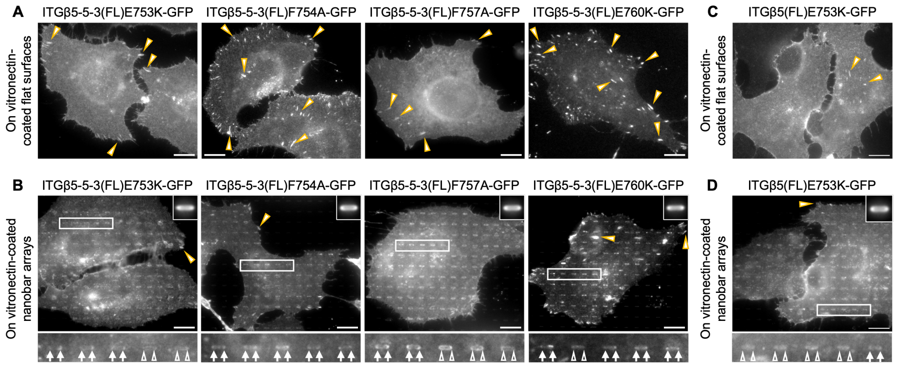


**Supplementary Figure 4. Mutation of the key residues in the membrane-proximal region of ITGβ5-5-3 reduces focal adhesion formation but increases curvature preference (Related to Figure 2).** **(A-B)** Representative images of ITGβ5-5-3 MP talin-binding mutants, including ITGβ5-5-3(E753K), ITGβ5-5-3(F754A), ITGβ5-5-3(F757A), and ITGβ5-5-3(E760K), on flat substrates (A) or nanobar arrays (B). These ITGβ5-5-3 mutants form much reduced focal adhesions compared to the wild-type ITGβ5-5-3. **(C-D)** Representative images of ITGβ5(E753K) on flat substrates (C) or nanobar arrays (D). ITGβ5(E753K) forms considerably fewer focal adhesions than the wild-type ITGβ5. Scale bar: 10 μm for all the cell images. White arrows indicate enrichments at nanobar ends, white empty arrowheads indicate no preferential enrichment at nanobar ends, while yellow arrowheads indicate focal adhesions. In (B) and (D), the averaged nanobar images are shown on the top-right corners.

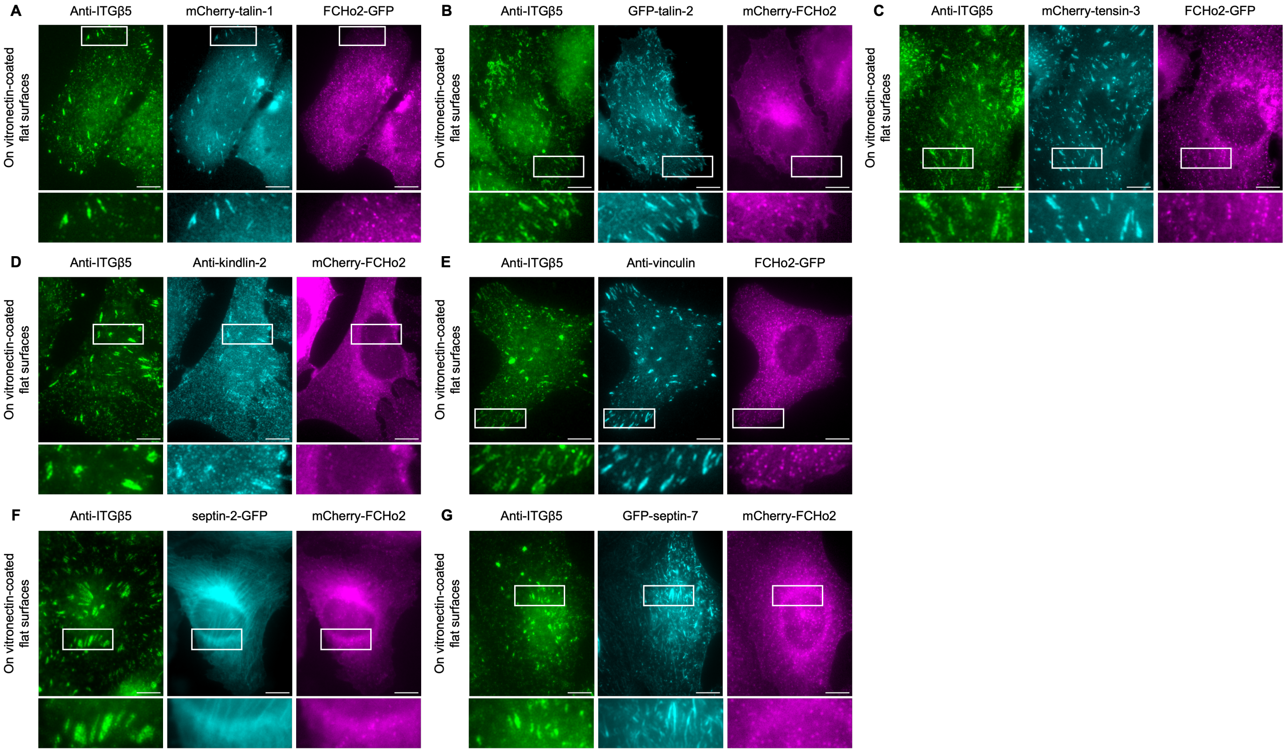


**Supplementary Figure 5. ITGβ5 colocalizes with a number of cytoskeletal adaptors in focal adhesions formed on flat substrates (Related to Figure 3). (A-E)** mCherry-talin-1 (A), GFP-talin-2 (B), mCherry-tensin-3 (C), anti-kindlin-2 (D), anti-vinculin (E), but not FCHo2 (mCherry- or GFP-tagged), all colocalize with anti-ITGβ5 in focal adhesion architectures formed on flat substrates. **(F-G)** On flat substrates, anti-ITGβ5 signals are exclusively localized to focal adhesions, whereas septin-2-GFP (F) and GFP-septin-7 (G) form filamentous structures. mCherry-FCHo2 is present in neither architecture. Scale bar: 10 μm for all the images.

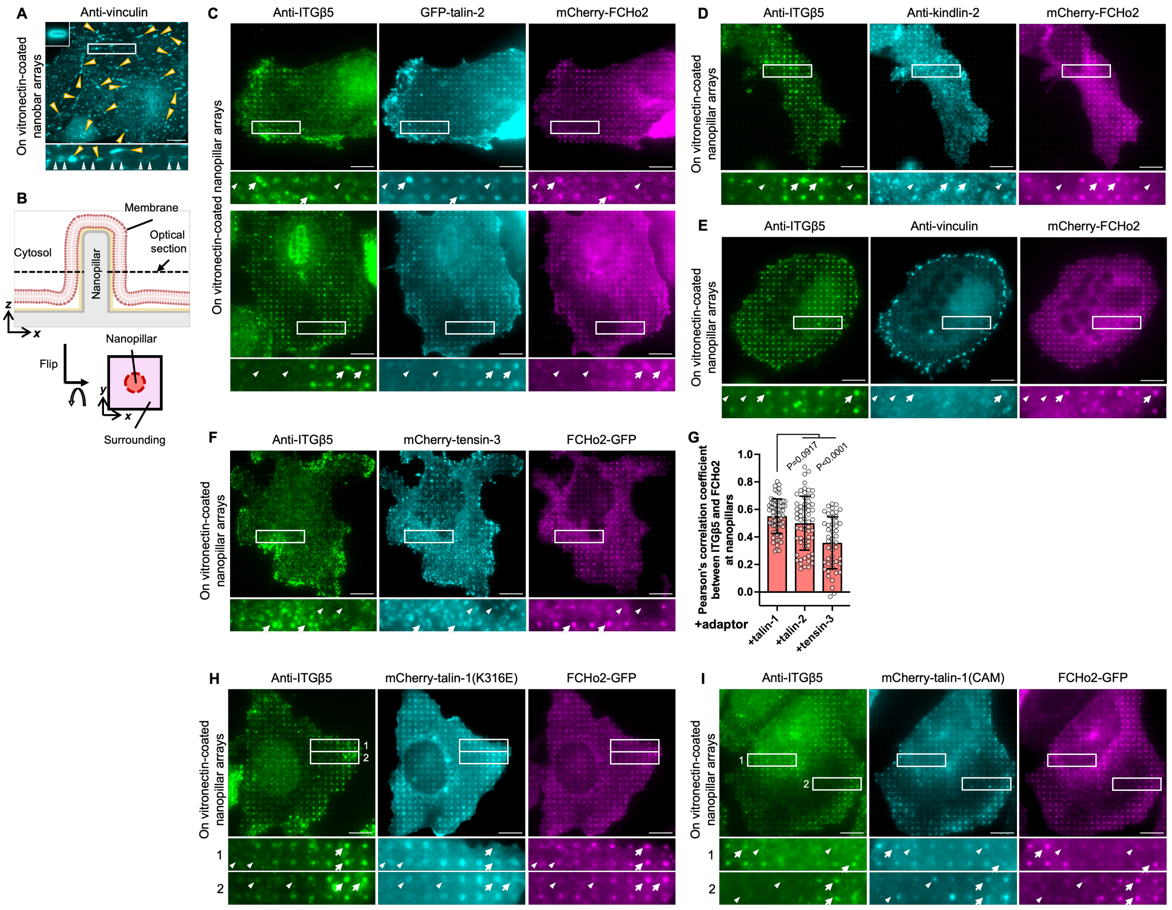


**Supplementary Figure 6. Talin-1, but not kindlin, tensin, or septin, is the primary mechanosensitive component in curved adhesions (Related to Figure 3).** **(A)** Representative images of anti-vinculin on nanobar substrates. Vinculin exhibits no apparent curvature preference (white empty arrowheads) while is exclusively enriched in focal adhesion patches formed on the flat regions of nanobar substrates (yellow arrowheads). **(B)** Schematic illustrations of a cell interfacing with a single nanopillar which induces cylindrical curvatures along its height. **(C)** GFP-talin-2 moderately accumulates and colocalizes with anti-ITGβ5 and mCherry-FCHo2 in curved adhesions formed at nanopillar locations. **(D-E)** Neither kindlin-2 (D) nor vinculin (E) is involved in curved adhesions, as seen by their modest-to-low accumulations and low spatial correlations with anti-ITGβ5 and mCherry-FCHo2, at nanopillars. **(F)** In some cells, overexpression of mCherry-tensin-3 attenuates curved adhesion formation, as reflected by reduced ITGβ5 accumulations at nanopillars. **(G)** Quantifications of Pearson's correlation coefficients between anti-ITGβ5 and FCHo2 (GFP- or mCherry-tagged) when co-expressed with mCherry-talin-1, GFP-talin-2, or mCherry-tensin-3. **(H)** mCherry-talin-1(K316E) shows a reduced colocalization with anti-ITGβ5 at nanopillars. **(I)** mCherry-talin-1(CAM) displays low spatial correlations with anti-ITGβ5 and FCHo2-GFP at nanopillars. Scale bar: 10 μm for all the cell images. In (C)-(F) and (H)-(I), white arrows indicate high-intensity correlations, while white triangles indicate low-intensity correlations at nanopillars. Welch's t tests (unpaired, two-tailed, not assuming equal variance) are applied for statistical analyses of curvature-related measurements. Error bars represent standard deviations. Each data point represents one cell.

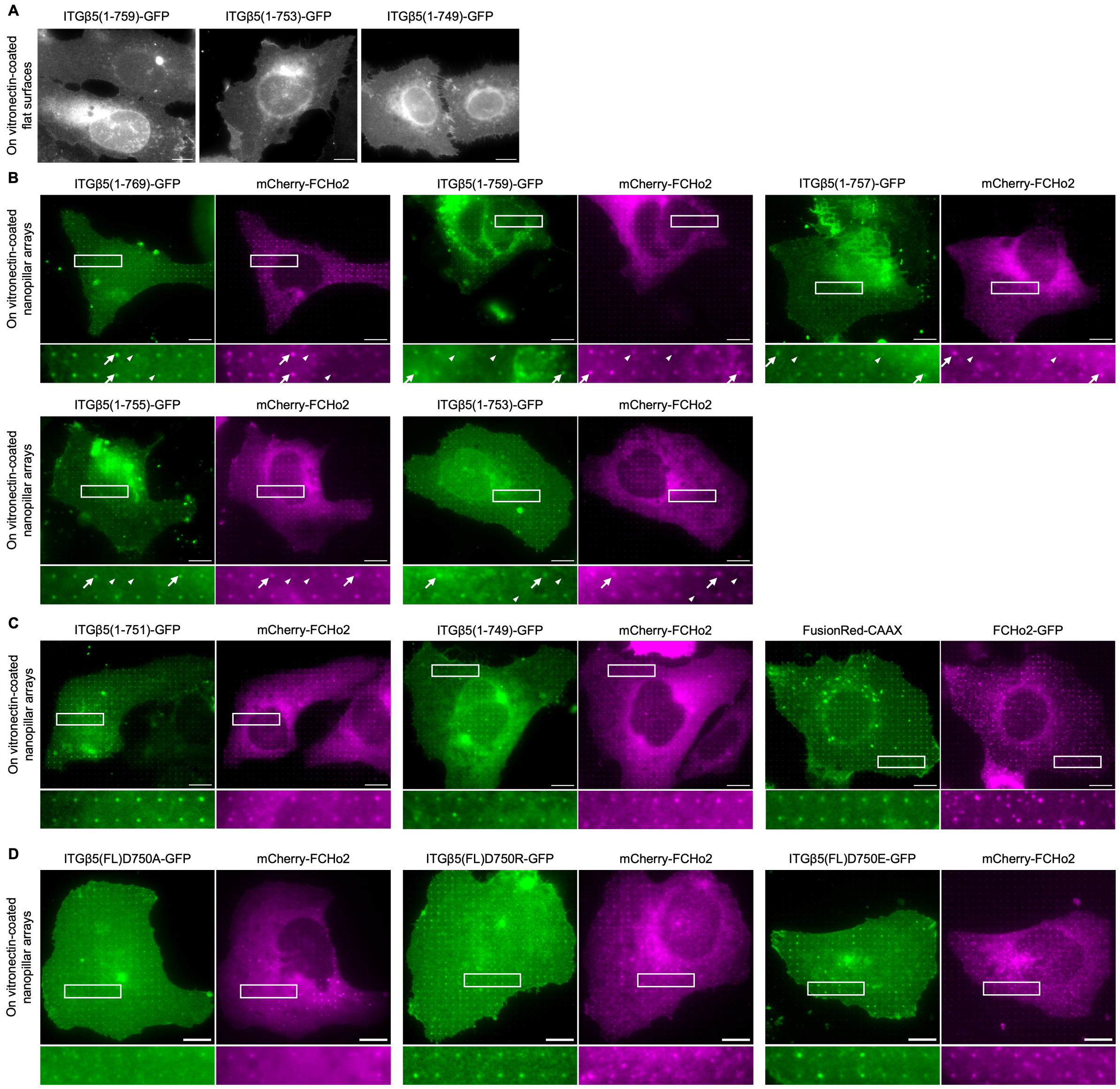


**Supplementary Figure 7. Co-expression of mCherry-FCHo2 and ITGβ5 truncation variants or D750 mutants on vitronectin-coated nanopillar arrays (Related to Figure 4).** **(A)** Representative images of three ITGβ5 truncation variants, including ITGβ5(1-759), ITGβ5(1-753), and ITGβ5(1-749), on flat substrates. **(B)** On nanopillar substrates, ITGβ5(1-769)-GFP, ITGβ5(1-759)-GFP, ITGβ5(1-757)-GFP, ITGβ5(1-755)-GFP, and ITGβ5(1-753)-GFP are all spatially correlated with mCherry-FCHo2 at nanopillar locations. White arrows indicate high-intensity correlations and white triangles indicate low-intensity correlations at nanopillars. **(C)** Neither ITGβ5(1-751)-GFP nor ITGβ5(1-749)-GFP exhibits an apparent spatial correlation with mCherry-FCHo2 at nanopillars, resembling the membrane marker FusionRed-CAAX. **(D)** mCherry-FCHo2 exhibits no apparent spatial correlations with ITGβ5(D750A)-GFP, ITGβ5(D750R)-GFP, or ITGβ5(D750E)-GFP, at nanopillars. Scale bar: 10 μm for all the images.

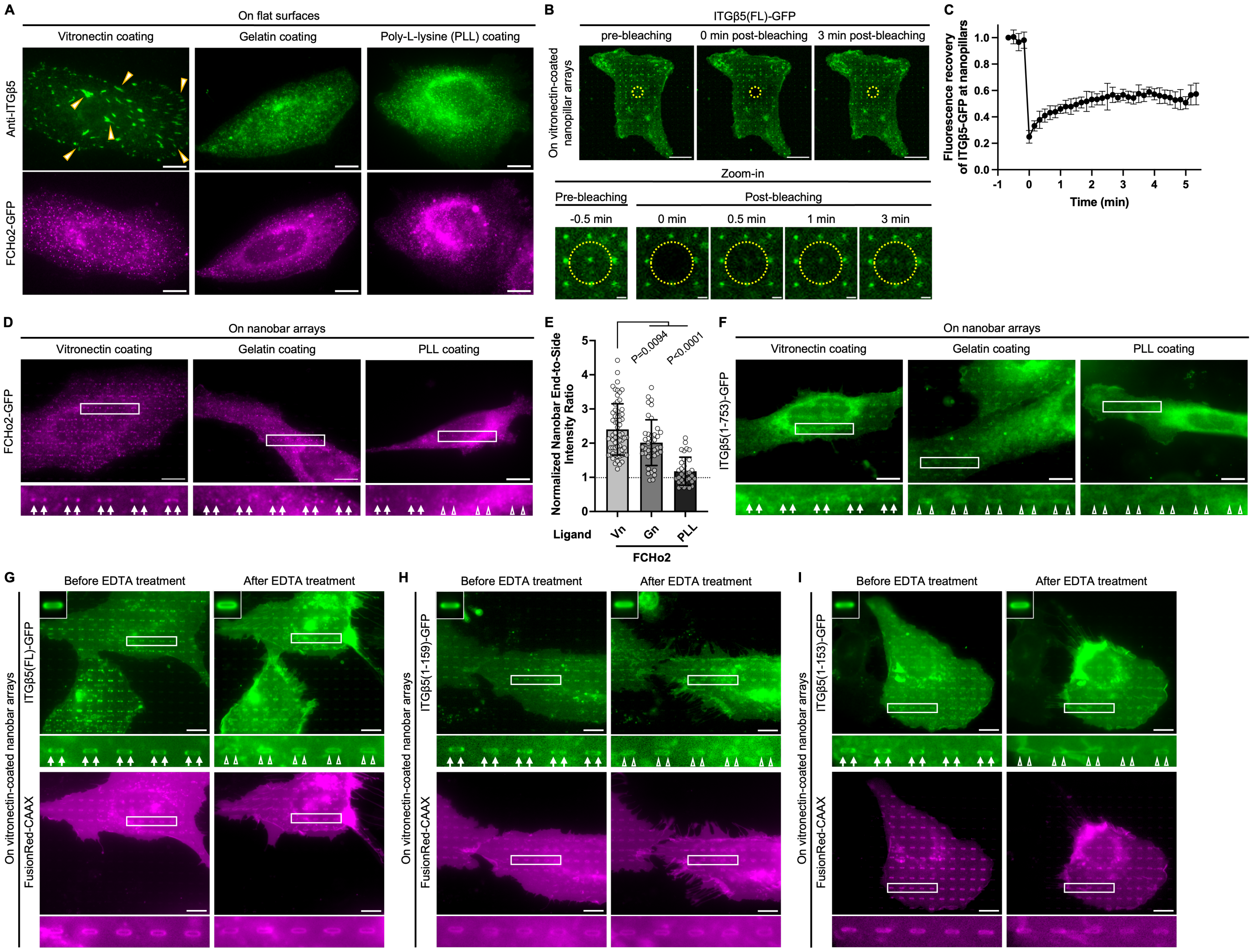


**Supplementary Figure 8. Curved adhesions involve active integrin αvβ5 (Related to Figure 5).** **(A)** ITGβ5 forms extensive focal adhesions on vitronectin-coated flat substrates (left), but not on gelatin- (middle) or PLL-coated (right) ones. Yellow arrowheads indicate focal adhesions. **(B)** To perform fluorescence recovery after photobleaching (FRAP), a region of interest (yellow dashed circle) was bleached by a 488-nm laser. ITGβ5-GFP signals at nanopillars were only partially recovered within 3 minutes post photobleaching, indicating that ITGβ5-mediated curved adhesions are stable. Scale bar: 10 µm for the larger-field images, 1 µm for all the zoom-in images. **(C)** Time course of FRAP signals of ITGβ5-GFP at nanopillars. **(D)** FCHo2-GFP displays a strong preference for nanobar ends on vitronectin-coated substrates. However, the preferential enrichments of FCHo2-GFP at nanobar ends are reduced when substrates are coated with gelatin or poly-L-lysine. **(E)** Quantifications of the normalized nanobar end-to-side intensity ratios of FCHo2-GFP on nanobar substrates coated with different ligands. Vn: vitronectin; Gn: gelatin; PLL: poly-L-lysine. The dashed line indicates a value of 1. **(F)** ITGβ5(1-753)-GFP displays a clear preference for nanobar ends only when substrates are coated with vitronectin. **(G-I)** After 5-min incubation, EDTA dramatically reduces accumulations of the full-length ITGβ5-GFP (G), ITGβ5(1-759)-GFP (H), and ITGβ5(1-753)-GFP (I) at nanobar ends on vitronectin-coated substrates. Within the time period, cell membranes remain wrapping around nanobars. Image sets of ITGβ5(1-759)-GFP are also shown in Figure 5E. Scale bar: 10 µm for all the images except for some in (B). White arrows indicate enrichments at nanobar ends, while white empty arrowheads indicate no preferential enrichment at nanobar ends. In (G)-(I), the averaged nanobar images are shown on the top-left corners. Welch's t tests (unpaired, two-tailed, not assuming equal variance) are applied for statistical analyses of curvature-related measurements. Error bars represent standard deviations. Each data point represents one cell.

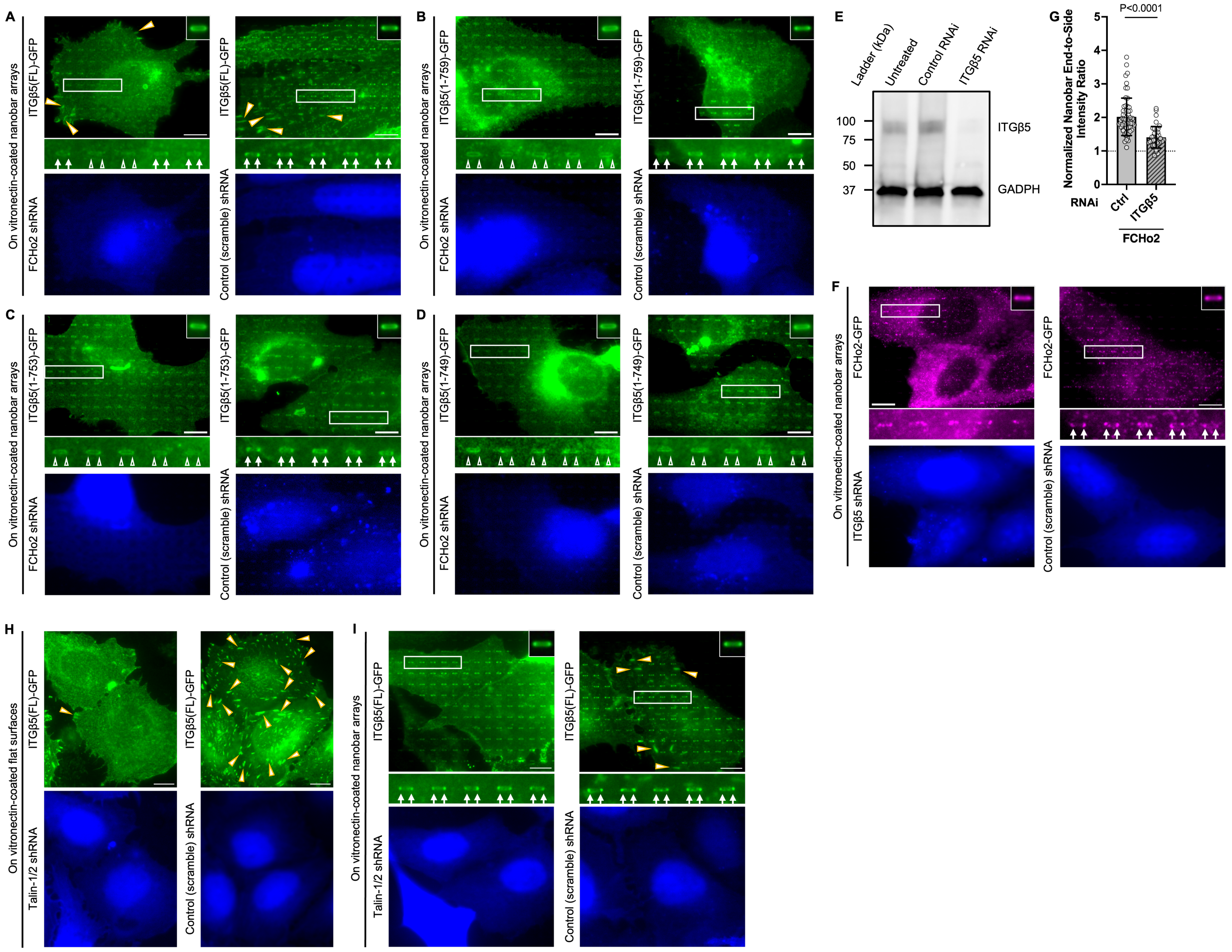


**Supplementary Figure 9. Curvature enrichments of ITGβ5-GFP (and its truncation variants) are significantly diminished upon FCHo2 knockdown, but retained upon double knockdown of talin-1 and talin-2 (Related to Figure 5).** **(A-C)** Preferential accumulations of the full-length ITGβ5-GFP (A), ITGβ5(1-759)-GFP (B), and ITGβ5(1-753)-GFP (C) at nanobar ends are eliminated upon FCHo2 knockdown by shRNA (left images), but not by treatment with control (scramble) shRNA (right images). Image sets of ITGβ5(1-759)-GFP are also shown in Figure 5H. **(D)** ITGβ5(1-749)-GFP exhibits no apparent curvature preference regardless of the shRNA treatment. **(E)** Western blots confirm the efficient knockdown of ITGβ5 by shRNA. **(F)** FCHo2-GFP accumulations at nanobar ends are significantly reduced upon ITGβ5 knockdown (left images), but not by treatment with control shRNA (right images). **(G)** Quantifications of the normalized nanobar end-to-side intensity ratio of FCHo2-GFP upon treatment with ITGβ5 shRNA or control shRNA. The dashed line indicates a value of 1. **(H)** ITGβ5-marked focal adhesions are drastically diminished upon double knockdown of talin-1 and talin-2 (left images), but retained upon treatment with control shRNA (right images). Image sets of ITGβ5(FL)-GFP are also shown in Figure 5K. **(I)** Curved adhesions, illustrated by the ITGβ5-GFP enrichments at nanobar ends, remain intact upon double knockdown of talin-1 and talin-2. Image sets of ITGβ5(FL)-GFP are also shown in Figure 5M. Scale bar: 10 μm for all the images. White arrows indicate enrichments at nanobar ends, white empty arrowheads indicate no preferential enrichment at nanobar ends, while yellow arrowheads indicate focal adhesions. In (A)-(D), (F), and (I), the averaged nanobar images are shown on either the top-right corners. Welch's t tests (unpaired, two-tailed, not assuming equal variance) are applied for statistical analyses of curvature-related measurements. Error bars represent standard deviations. Each data point represents one cell.

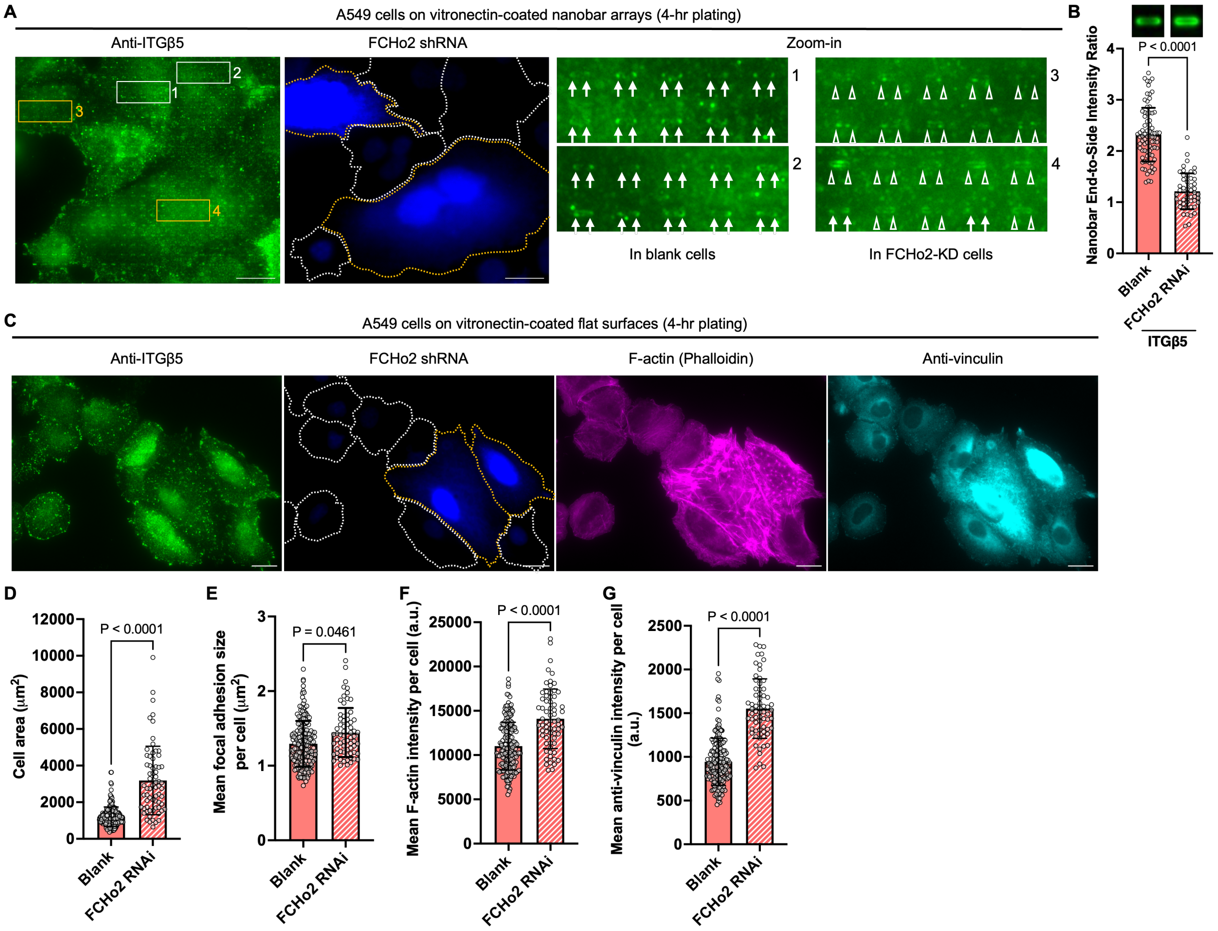


**Supplementary Figure 10. Depletion of FCHo2 drastically enhances focal adhesion formation in A549 cells (Related to Figure 5).** **(A)** Preferential accumulations of the endogenous ITGβ5 at nanobar ends are eliminated in FCHo2-depleted A549 cells (outlined by yellow dotted lines), but not in blank or mildly-depleted ones (outlined by white dotted lines). White arrows indicate enrichments at nanobar ends, white empty arrowheads indicate no preferential enrichment at nanobar ends, while yellow arrowheads indicate focal adhesions. **(B)** Quantifications of the nanobar end-to-side intensity ratio of anti-ITGβ5-GFP in blank or FCHo2-depleted A549 cells. The averaged nanobar images are shown on the top. **(C)** On flat substrates, FCHo2-depleted A549 cells are visibly larger than blank or mildly-depleted ones. In addition, F-actin and anti-viculin intensities are significantly elevated in FCHo2-depleted A549 cells. **(D-G)** Quantifications of cell area (D), mean focal adhesion area per cell (E), mean F-actin intensity per cell (F), and mean anti-vinculin intensity per cell (G) in blank or FCHo2-depleted A549 cells. Scale bar: 20 μm for all the cell images. Welch's t tests (unpaired, two-tailed, not assuming equal variance) are applied for statistical analyses of curvature-related measurements (i.e. normalized nanobar end-to-side intensity ratios) and protein fluorescence intensity quantifications, while Kolmogorov-Smirnov test was used for statistical analyses of focal adhesions and cell area. Error bars represent standard deviations. Each data point represents one cell. Error bars represent standard deviations. Each data point represents one cell.

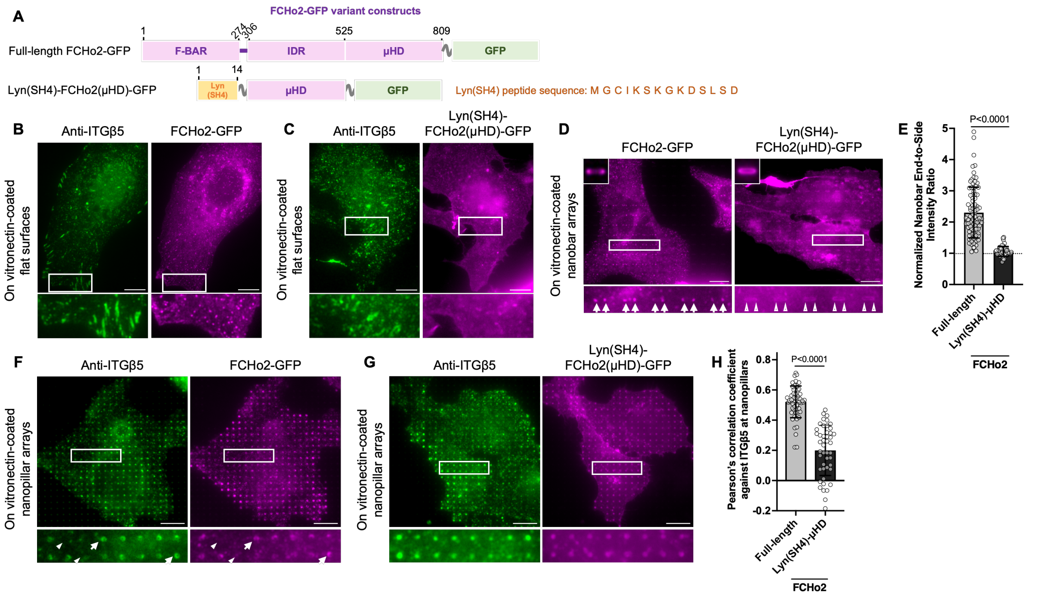


**Supplementary Figure 11. Both the F-BAR domain and μHD of FCHo2 are required for curvature-dependent recruitment of integrin αvβ5 curved adhesions (Related to Figure 5).** **(A)** Domain structures of two FCHo2-GFP variants. IDR: intrinsically disordered region; μHD: microhomology domain. **(B-C)** Neither the wild-type FCHo2-GFP (B) nor Lyn(SH4)-FCHo2(μHD)-GFP (C) is localized to ITGβ5-marked focal adhesion patches on flat substrates. **(D)** On nanobar arrays, FCHo2-GFP displays a clear curvature preference, while Lyn(SH4)-FCHo2(μHD)-GFP behaves like a membrane marker by uniformly wrapping around nanobars. White arrows indicate enrichments at nanobar ends, while white empty arrowheads indicate no preferential enrichment at nanobar ends. The averaged nanobar images are shown on the top-left corners. **(E)** Quantifications of the normalized nanobar end-to-side intensity ratio of two FCHo2-GFP variants. The dashed line indicates a value of 1. **(F-G)** On nanopillar substrates, FCHo2-GFP (F), but not Lyn(SH4)-FCHo2(μHD)-GFP (G), spatially correlates with the endogenous ITGβ5 at nanopillars. White arrows indicate high-intensity correlations, while white triangles indicate low-intensity correlations at nanopillars. **(H)** Quantifications of Pearson's correlation coefficients between anti-ITGβ5 and two FCHo2-GFP variants at nanopillars. Scale bar: 10 μm for all the images. Welch's t tests (unpaired, two-tailed, not assuming equal variance) are applied for statistical analyses of curvature-related measurements. Error bars represent standard deviations. Each data point represents one cell.

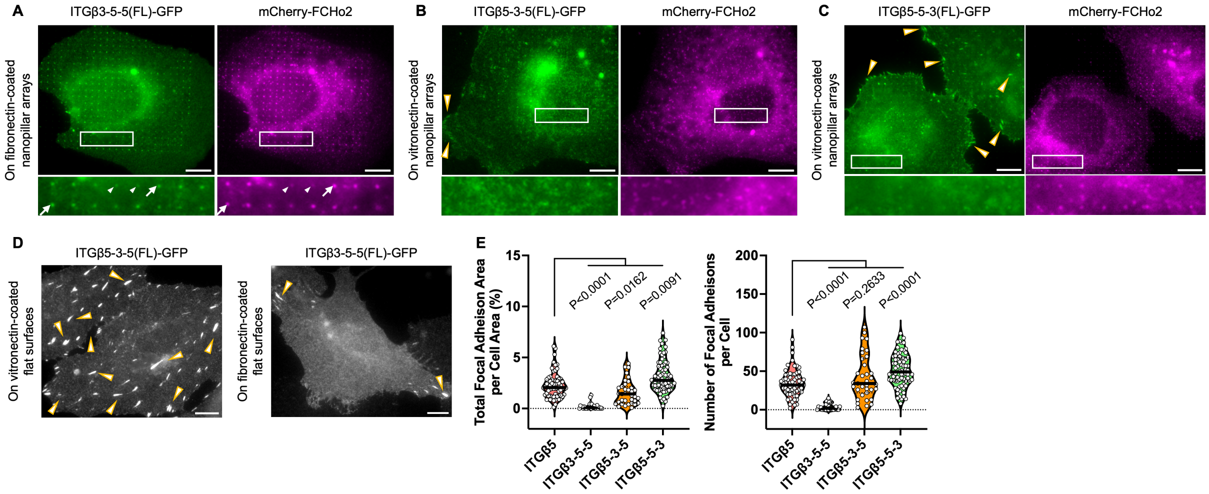


**Supplementary Figure 12. ITGβ3-5-5-GFP, but neither ITGβ5-3-5-GFP nor ITGβ5-5-3-GFP, spatially correlates with mCherry-FCHo2 at nanopillars (Related to Figure 6).** **(A-C)** mCherry-FCHo2 accumulates and spatially correlates with ITGβ3-5-5(FL)-GFP (A), but with neither ITGβ5-3-5(FL)-GFP (B) nor ITGβ5-5-3(FL)-GFP (C), at nanopillars. Among the three chimeric variants, only ITGβ3-5-5-GFP demonstrates curvature sensitivity and clearly accumulates in curved adhesions formed at nanopillars. White arrows indicate high-intensity correlations, while white triangles indicate low-intensity correlations at nanopillars. **(D)** Representative images of ITGβ5-3-5-GFP (left) and ITGβ3-5-5-GFP (right) on flat substrates. ITGβ5-3-5 forms significantly more focal adhesion patches than ITGβ3-5-5. **(E)** Quantifications of the focal adhesion area percentage (left) and the number of focal adhesions per cell (right) of four integrin chimeric variants. Scale bar: 10 μm for all the images. Yellow arrowheads indicate focal adhesions. Kolmogorov-Smirnov test was used for statistical analyses of focal adhesions. Each data point represents one cell.

| Target protein | Source | Identifier | Target sequence | KD efficiency* |
|---|---|---|---|---|
| Integrin β5 | Sigma-Aldrich | TRCN0000296116 | GGATCAGCCTGAGGATCTTAA | 82% |
| | | TRCN0000296117 | AGCTTGTTGTCCCAATGAAAT | 85% |
| | | TRCN0000289121 | CTGAGGGCAAACCTTGTCAAA | 84% |
| FCHo2 | | TRCN0000167218 | GCTACAGTATTAAACCAGAAA | 88% |
| | | TRCN0000167925 | CCAAAGCTTACTTCAGGCAAA | 75% |
| Talin-1 | | TRCN0000299020 | GCCTCAGATAATCTGGTGAAA | 95% |
| | | TRCN0000299022 | CCCAGAGTATTAACGCTCCAA | 93% |
| | | TRCN0000299091 | GCAGTGAAAGATGTAGCCAAA | 75% |
| Talin-2 | | TRCN0000436291 | GACGAATCCAAACACGAAATC | N.A. |
| | | TRCN0000439801 | ACGATGCGTGTCGAGTCATTC | N.A. |
| Scramble (Control) | Addgene | #1864 | CCTAAGGTTAAGTCGCCCTCG | N.A. |

**Supplementary Table 1. Sequence and knockdown efficiency of shRNA used in this work.** *Knockdown efficiencies are obtained from Millipore Sigma "Predesigned shRNA" database.

| **Name** | **Supplier** | **Clone** | **Identifier** | **Dilution factor** |
|---|---|---|---|---|
| Primary antibodies | | | | |
| Rabbit anti-integrin β5 antibody | Cell signaling | D24A5 | #3629S | 1:500 (for IF)<br>1:2000 (for WB) |
| Mouse anti-vinculin antibody | Sigma-Aldrich | hVIN-1 | #V9131 | 1:500 (for IF) |
| Mouse anti-kindlin-2 antibody | Antibodies.com | 3A3 | #A277536 | 1:500 (for IF) |
| Rabbit anti-FCHo2 antibody | Novus Biologicals | Polyclonal | #NBP2-32694 | 1:1500 (for WB) |
| Rabbit anti-talin-1/2 antibody | abcam | 8D4 | ab11188 | 1:2000 (for WB) |
| Rabbit anti-GAPDH antibody | Cell signaling | 14C10 | #2118 | 1:5000 (for WB) |
| Rabbit anti-β-tubulin antibody | Cell signaling | D2N5G | #15115 | 1:5000 (for WB) |
| Secondary antibodies | | | | |
| Goat anti-Mouse IgG (H+L) Highly Cross-Adsorbed Secondary Anti-body, Alexa Fluor™ 488 | Invitrogen | N.A. | #A-32723 | 1:500 (for IF) |
| Goat anti-Mouse IgG (H+L) Highly Cross-Adsorbed Secondary Anti-body, Alexa Fluor™ 594 | Invitrogen | N.A. | #A-11032 | 1:500 (for IF) |
| Goat anti-Rabbit IgG (H+L) Highly Cross-Adsorbed Secondary Anti-body, Alexa Fluor™ Plus 647 | Invitrogen | N.A. | #A-32733 | 1:500 (for IF) |
| HRP-linked goat anti-Rabbit IgG (H+L) antibody | Cell signaling | N.A. | #7074 | 1:5000 (for WB) |
| HRP-linked goat anti-Mouse IgG (H+L) antibody | Cell signaling | N.A. | #7076 | 1:5000 (for WB) |

**Supplementary Table 2. Antibodies used in this work.**
